# Nuclear RNAi Modulates Influenza A Virus Infectivity By Downregulating Type-I Interferon Response

**DOI:** 10.1101/2024.03.07.583365

**Authors:** Hsiang-Chi Huang, Iwona Nowak, Vivian Lobo, Danica F. Besavilla, Karin Schön, Jakub O. Westholm, Carola Fernandez, Angana A.H. Patel, Clotilde Wiel, Volkan I. Sayin, Dimitrios G. Anastasakis, Davide Angeletti, Aishe A. Sarshad

## Abstract

The role of Argonaute (AGO) proteins and the RNA interference (RNAi) machinery in mammalian antiviral response has been debated. Therefore, we set out to investigate how mammalian RNAi impacts influenza A virus (IAV) infection. We reveal that IAV infection triggers nuclear accumulation of AGO2, which is directly facilitated by p53 activation. Mechanistically, we show that IAV induces nuclear AGO2 targeting of TRIM71, a proposed AGO2 E3 ligase, and type-I interferon-pathway genes for silencing. Accordingly, *Tp53^-/-^* mice do not accumulate nuclear AGO2 and demonstrate decreased susceptibility to IAV infection. Hence, the RNAi machinery is highjacked by the virus to evade the immune system and support viral replication. Furthermore, the FDA approved drug arsenic trioxide, which prevents p53 tetramerization and nuclear translocation, increases interferon response and decreases viral replication *in vitro* and in a mouse model *in vivo*. Our data indicates that targeting the AGO2:p53-mediated silencing of innate immunity may offer a promising strategy to mitigate viral infections.

## INTRODUCTION

Argonaute (AGO) proteins have a central role in RNA interference (RNAi), where they are guided by endogenous miRNA or exogenous siRNA to recognize (partially) complementary sequences on target RNAs ^1^. Together, AGO-small RNA (smRNA) constitutes the core of the RNA-induced silencing complex (RISC), a multiprotein complex that deregulates RNA transcripts, resulting in target destabilization ^1^. RNAi has a well-established role in antiviral defense in certain eukaryotes, including plants, insects and nematodes ^2,3^. However, in mammals, the role of RNAi as an antiviral defense mechanism is more controversial: indeed, while some studies described antiviral RNAi functions ^4,5^, many others reported a lack of evidence for direct antiviral activity ^6^. What has not been extensively investigated is whether RNAi could be hijacked by viruses to their advantage. Sparse evidence suggests that loss of RNAi after viral infection may indeed decrease viral titers with concomitant increased expression of antiviral genes ^6–8^. The overall picture and detailed mechanisms of action remain, however, elusive.

Upon infection with influenza A virus (IAV), the innate immune system serves as the body’s primary defense, swiftly initiating a response to combat invading pathogens ^9^. A crucial facet of the innate immune response to IAV is the synthesis of type-I interferons (IFN-Is) by infected cells. These interferons subsequently activate neighboring cells to produce antiviral proteins, which are essential in curtailing the severity and duration of infections ^10^. Viral RNAs, in their single-strand or intermediate format, can be recognized by the Toll-like receptors TLR3 and TLR7 to activate IFN transcription via NF-kβ, IRF-3 or IRF-7 ^9^. Collectively, viral RNAs therefore activate the immune system by different mechanisms, which all lead to interferon production. Additionally, the tripartite motif (TRIM) family of proteins modulates the production, signaling, and effector functions of IFN-Is, thereby influencing immune responses and host defense against pathogens. Notably, TRIM25, TRIM56, and TRIM71 exert positive regulatory roles in the IFN-I pathway ^11^.

From the IAV side, the multifunctional nonstructural protein 1 (NS1) of IAV serves as an RNA-binding protein, facilitating mRNA export from the nucleus. Moreover, unrelated to the nuclear export function, NS1 can re-enter the cell nucleus and has a pivotal role in antagonizing host immune responses and facilitating pathogenesis ^12–15^. Furthermore, NS1 has been shown to inhibit TRIM25 oligomerization, thus suppressing RIG-I-mediated IFN production ^16^. Therefore, NS1 is the key player in IAV innate immune evasion, with multiple functions, including the ability to block RISC ribosylation ^7^. It has been shown that ribosylation of the RISC complex leads to the shutdown of RNAi mechanisms. This is intriguing because it suggests that it is in the virus’ best interest to maintain RNAi function. However, while it was shown that two specific miRNAs could inhibit interferon-stimulated genes (ISGs) in IFN-treated cells, there are still open questions regarding global, mechanistic functions of RNAi after viral infections.

Adding to the complexity of viral regulation of RISC function is the identification of nuclear RNAi. In metazoans, cytoplasmic RNAi processes are well documented ^17^, yet the core RISC components – AGOs and miRNAs – have been found in cell nuclei ^18–21^. Indeed, various mechanisms of stress-induced nuclear translocation of AGOs have been proposed, including DNA damage and viral infection ^22,23^. While cytoplasmic AGO2 is well known as a critical component of RISC and is involved in siRNA/miRNA-mediated gene suppression pathways ^24^, little is known about the specific functions of nuclear AGO2. This is particularly relevant to disentangle the controversies around the role of RNAi in mammals after viral infections.

To comprehensively address the role of RNAi following IAV infection, we combined *in vitro* and *in vivo* experimental models with fluorescence-based photoactivatable ribonucleoside-enhanced crosslinking and immunoprecipitation (fPAR-CLIP) to pinpoint AGO2 targets at nucleotide resolution. We demonstrate that upon IAV infection, NS1 mediates AGO2 translocation into the cell nucleus to silence TRIM71 and IFN-pathway-related genes, thereby increasing viral replication. Our data provide important mechanistic insights into previously underappreciated modalities of viral resistance.

## RESULTS

### Viral infection induces nuclear accumulation of AGO2

AGO2 may localize both in the cytoplasm and the nucleus of human cells ^25^. For instance, we previously demonstrated the absence of nuclear AGO2 in HEK293 cells ^21,25^. However, we observed a high degree of nuclear AGO2 in HEK293T cells, which are HEK293 cells transformed with SV40 large T (LT) antigen, the master regulator of polyomaviruses ^26^ (**Fig 1A**). Biochemical fractionation of HEK293T cells showed a near 50-50 distribution of AGO2 between the cytoplasmic and nuclear fractions (**Fig 1A**). Moreover, transient overexpression of SV40 LT antigen in HEK293 cells led to the translocation of AGO2 into the nucleus (**Fig 1B**), indicating the involvement of SV40 LT antigen in AGO2 nuclear accumulation. These observations prompted us to investigate whether nuclear translocation of AGO2 is shared by acute viral infections.

**Figure 1:**
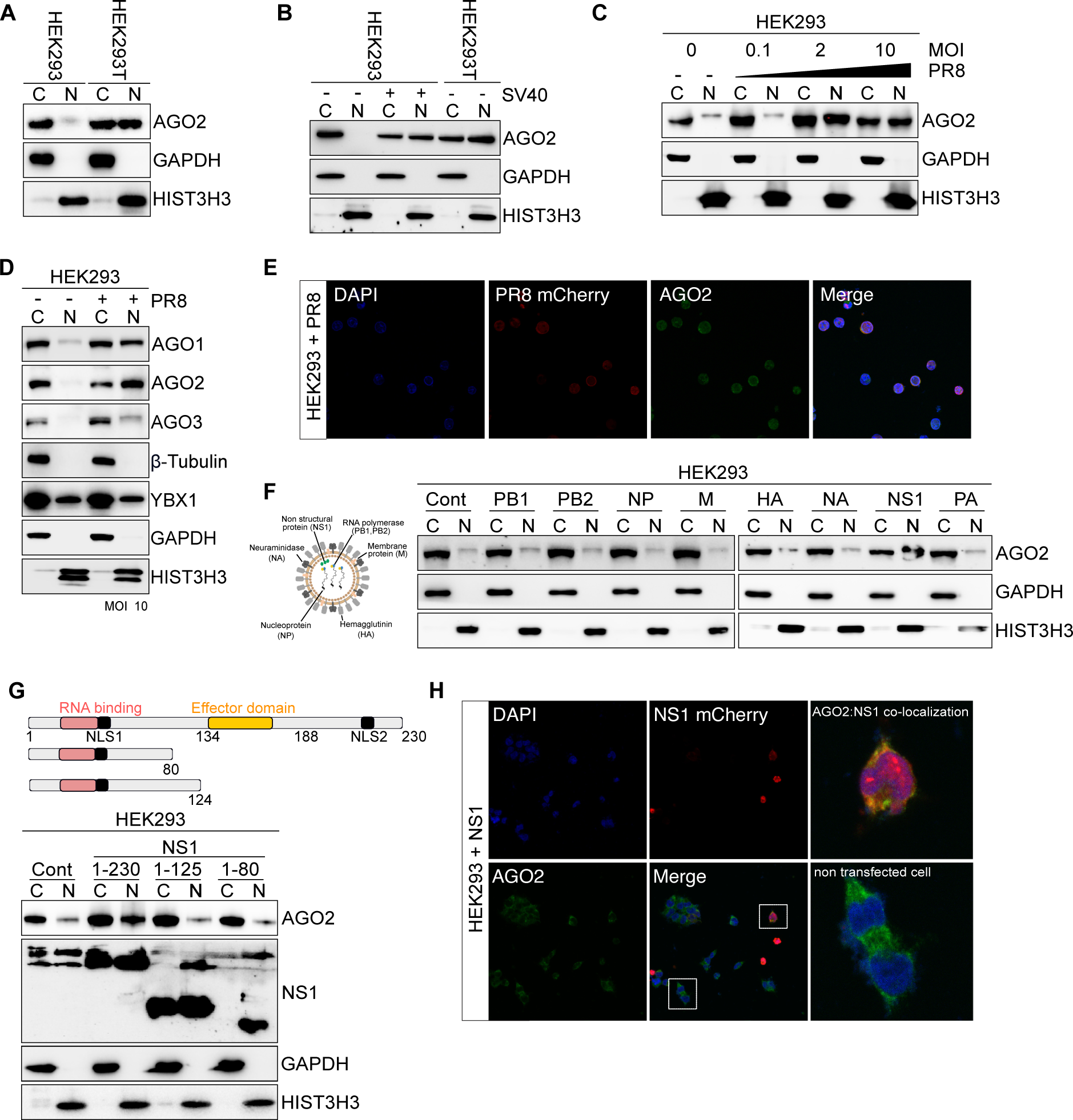
IAV virus NS1 induces AGO2 nuclear translocation. **(A)** Representative AGO2 immunoblots from cytoplasmic (C) and nuclear (N) lysates in HEK293 and HEK293T cells. GAPDH served as cytoplasmic marker and HIST3H3 as nuclear marker. n=3 **(B)** Representative AGO2 immunoblots from cytoplasmic (C) and nuclear (N) lysates in HEK293 cells transfected with plasmid expressing SV40 Large T antigen. HEK293T was used as a positive control. GAPDH served as cytoplasmic marker and HIST3H3 as nuclear marker. n=3 **(C)** Representative AGO2 immunoblots from cytoplasmic (C) and nuclear (N) lysates in HEK293 cells infected with PR8 virus at MOI 0.1; 2; 10 for 16 hours. GAPDH served as cytoplasmic marker and HIST3H3 as nuclear marker. n=3 **(D)** Representative AGO1, AGO2 and AGO3 immunoblots from cytoplasmic (C) and nuclear (N) lysates in HEK293 cells infected with PR8 virus at MOI 10 for 16 hours. GAPDH and β-Tubulin served as cytoplasmic marker and HIST3H3 as nuclear marker. YBX1 served as a control for shuttling protein. n=3 **(E)** Immunofluorescence images of AGO2 and PR8-mCherry in HEK293 infected with PR8-NS1-mCherry virus at MOI 10 for 16 hours. DAPI stained for DNA. **(F)** Representative AGO2 immunoblots from cytoplasmic (C) and nuclear (N) lysates in HEK293 cells transfected with PB1, PB2, NP, M, HA, NS1, and PA expressing plasmids for 2 days. GAPDH served as cytoplasmic marker and HIST3H3 as nuclear marker. n=3 **(G)** Representative AGO2 and NS1 immunoblots from cytoplasmic (C) and nuclear (N) lysates in HEK293 cells transfected with WT (1-230) and deletion mutant NS1 (1-80 and 1-124) expressing plasmid for 2 days. GAPDH served as cytoplasmic marker and HIST3H3 as nuclear marker. n=3 **(H)** Immunofluorescence images of AGO2 and NS1-mCherry in HEK293 infected with NS1-mCherry virus at MOI 10 for 16 hours. DAPI stained for DNA. Upper box highlights a cell where AGO2 and NS1-mCherry are colocalized in the nucleus, the lower box highlights cells that were not transfected with NS1 and AGO2 remains cytoplasmic.

To address this, we took advantage of PR8 strain of IAV as a model system. PR8 is favored in IAV research for its well-defined genetics, ease of modification, availability of mutants and ability to infect mice ^27^. HEK293 cells were infected with PR8-IAV at several multiplicity of infection (MOI) for 16 hours, and biochemical fractionation experiments were performed (**Fig 1C**). We observed robust nuclear translocation of AGO2 upon infection with PR8-IAV, particularly at MOI 2 or higher (**Fig 1C**). MOI lower than 2 did not yield any nuclear translocation of AGO2 after 16 hours of infection, suggesting that MOI < 2 may need longer time of infection. However, IAV infection must be performed in serum-free media which, by itself, induced cellular stress and AGO2 nuclear translocation (**Sup Fig 1A**). Therefore, all subsequent experiments were conducted at 16 hours of viral infection at MOI 2 or 10. Under these conditons, we also observed the nuclear translocation of AGO1 and AGO3 proteins upon viral infection (**Fig 1D**), suggesting that the nuclear accumulation of RNAi factors is a general phenomenon during IAV infection. To visualize the nuclear accumulation of AGO2 upon viral infection, we utilized a fluorescently tagged PR8 virus strain, which has mCherry inserted within the NS1 protein ^28^, and performed confocal microscopy (**Fig 1E, Sup Fig 1B**).

IAV consists of eight gene segments that encode up to 17 viral proteins ^29^. To identify the specific gene segment responsible for AGO2 nuclear accumulation, we transiently expressed each influenza protein in HEK293 cells and performed biochemical fractionation assays. Interestingly, we found that NS1 is the primary viral factor triggering AGO2 nuclear translocation (**Fig 1F**). NS1 has a N-terminal RNA binding domain, containing an NLS signal, and a C-terminal effector domain, interacting with cellular signals and regulating their functions ^30^. Furthermore, the first 113 amino acids are sufficient for normal RNA binding activity ^31^. The effector domain, comprising amino acid residues 86-205, is crucial for its function ^32^. Therefore we generated two NS1 truncated mutants and demonstrated that NS1-effector domain was indeed responsible for the nuclear translocation of AGO2 (**Fig 1G**). Furthermore, we observed nuclear co-localization of AGO2 with NS1 after transient expression of mCherry-tagged NS1 protein (**Fig 1H, Sup Fig 1C**). Collectively, these findings suggest that viral proteins, including SV40 LT antigen and IAV NS1, induce nuclear accumulation of AGO1-4.

### AGO2 interacts with p53 in the nucleus upon viral infection

Having established the nuclear translocation of AGO2, after IAV infection, we next wanted to identify potential interacting partners of AGO2 involved in its translocation or stabilization. Therefore, we retrieved a list of AGO2-associated proteins (**Sup Table 1**) from the Harmonizome database ^33^, and performed a protein-protein interaction network analysis by STRING (**Fig 2A**). STRING aggregates diverse data to map protein-protein interactions, enhancing understanding of molecular functions and cellular processes. We identified that AGO2 directly interacts with p53, as well as with other well-known components of the RISC complex (**Fig 2A**). Interestingly, SV40 LT antigen also possesses a p53 binding domain ^34^, Therefore, we wondered whether p53 may play a role in the nuclear translocation of AGO2 after IAV infection and whether p53 can also interact with the NS1 component of IAV.

**Figure 2:**
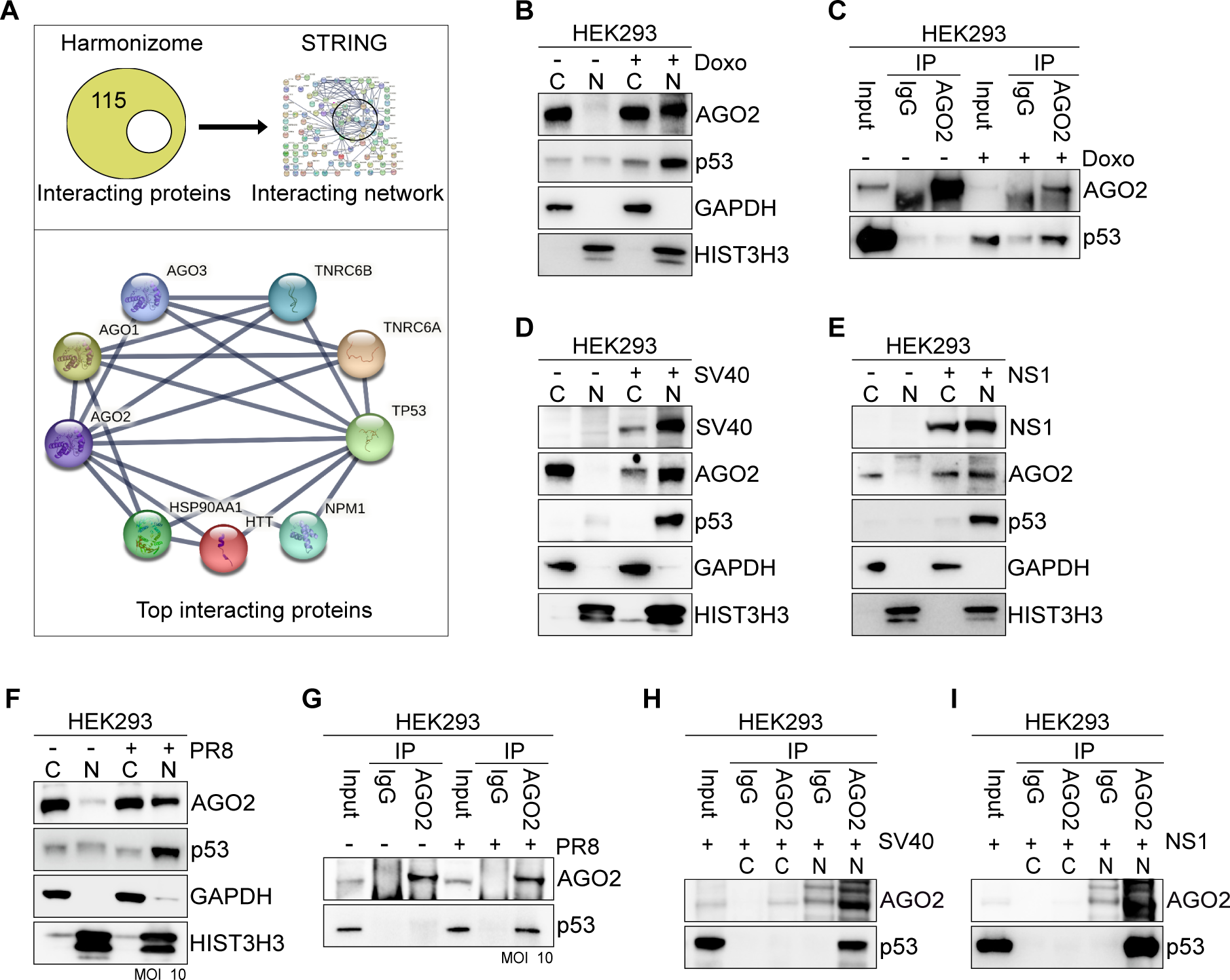
p53 associates with AGO2 in the nucleus. **(A)** STRING protein-protein interaction network of AGO2. **(B)** Representative AGO2 and p53 immunoblots from cytoplasmic (C) and nuclear (N) lysates in HEK293 cells treated with Doxorubicin (Doxo) for 24 hours. GAPDH served as a cytoplasmic marker and HIST3H3 served as nuclear marker. n=3 **(C)** AGO2 immunoprecipitation (IP) from HEK293 cells treated with Doxorubicin (Doxo) for 24 hours. Representative immunoblots of AGO2 and p53. n=3 **(D)** Representative AGO2, p53 and SV40 immunoblots from cytoplasmic (C) and nuclear (N) lysates in HEK293 cells transfected with SV40 large T antigen expressing plasmid. GAPDH served as a cytoplasmic marker and HIST3H3 served as nuclear marker. n=3 **(E)** same as in **(D)** except for immunoblots for NS1 in HEK293 cells transfected with NS1 mCherry expressing plasmid for 24 hours. n=3 **(F)** Representative AGO2 and p53 immunoblots from cytoplasmic (C) and nuclear (N) lysates in HEK293 cells infected with PR8 virus at MOI 10 for 16 hours. GAPDH served as cytoplasmic marker and HIST3H3 as nuclear marker. n=3 **(G)** AGO2 immunoprecipitation (IP) from HEK293 cells treated with PR8 virus at MOI 10 for 16 hours. Representative immunoblots of AGO2 and p53. n=3 **(H)** AGO2 immunoprecipitation (IP) from cytoplasmic and nuclear fractions in HEK293 cells transfected with SV40 large T antigen expressing plasmid for 24 hours. Representative immunoblots of AGO2 and p53. n=3 **(I)** same as in **(H)** except for HEK293 cells were transfected with WT NS1 expressing plasmid for 24 hours. n=3

To address these questions, we first examined the subcellular localization of p53 in HEK293 cells. We observed that p53 levels are low, yet ubiquitous, in HEK293 cells but, as reported, the majority of p53 translocated into the nucleus when cells were treated with doxorubicin, a DNA damage-inducing agent, for 24 hours (**Fig 2B**) ^35^. Consistent with our hypothesis, AGO2 also translocated into the nucleus in doxorubicin-treated cells (**Fig 2B**). Furthermore, by performing a co-immunoprecipitation experiment, we demonstrated a direct interaction between AGO2 and p53 upon doxorubicin treatment (**Fig 2C**). Interestingly, in lung cancer cells A549 (**Sup Fig 2A**), HEK293T (**Sup Fig 2B**), neuroblastomas cells SK-N-BE(2) (**Sup Fig 2C**) and breast cancer cell line MCF7 (**Sup Fig 2D**) AGO2 and p53 are ubiquitously expressed but interact exclusively within the nucleus (**Sup Fig 2A-D**).

Next, we investigated the subcellular localization of p53 in response to transient expression of SV40 LT antigen (**Fig 2D**) and NS1 (**Fig 2E and Sup Fig 2E**). Indeed, overexpression of both viral proteins induced substantial translocation of p53 into the nucleus, together with AGO2 (**Fig 2D,E**). Furthermore, in IAV-infected HEK293 cells, we observed not only nuclear AGO2 but also nuclear p53 accumulation (**Fig 2F**). Finally, to test whether AGO2 and p53 interact with each other also in response to viral infection, we co-immunoprecipitated AGO2 with p53. Indeed, we observed an interaction between AGO2 and p53 in virus-infected cells but not in control cells (**Fig 2G**). Similarly, we found that p53:AGO2 co-immunoprecipitated after transient expression of either SV40 LT (**Fig 2H**) or NS1 antigen (**Fig 2I**) but not NS1 mutant (**Sup Fig 2F)**.

Taken together, our findings suggest that both AGO2 and p53 translocate into the nucleus where they interact with each other in response to IAV infection. Furthermore, in cells, such as A549, which already have nuclear AGO2, p53 and AGO2 interact, indicating a potential functional interplay between these two proteins in the nucleus.

### The N-terminus of tetrameric p53 interacts with AGO2 and protects AGO2 from proteasomal degradation in the nucleus

We next wanted to examine how AGO2:p53 interact and what functional outcome the interaction may have in the nucleus. p53 is a 53 kDa protein consisting of an N-terminal transactivation domain, proline-rich domain, a core DNA binding domain and a C-terminal tetramerization and regulatory domain ^36^ (**Fig 3A**). The PIWI domain of AGO2 contains tandem tryptophan-binding pockets, which collectively form a region for interacting with TNRC6 or other tryptophan-rich cofactors ^37,38^. Interestingly, we observed the presence of three tryptophan residues (Trp23, Trp53, and Trp91) within the flexible N-terminal loop region of p53 (**Sup Fig 3A**), suggesting that the N-terminus may be involved in AGO2 binding. To investigate the possible interaction between the N-terminal region of p53 and AGO2, we generated two Flag-p53 mutants by removing amino acids 1-61 and 1-92 from the N-terminus (**Fig 3B**). Upon transient overexpression of these mutants in doxorubicin-treated HEK293 cells, we found that AGO2 exclusively interacts with full-length Flag-p53, while the interaction was abolished in both N-terminal mutants (**Fig 3C**).

**Figure 3:**
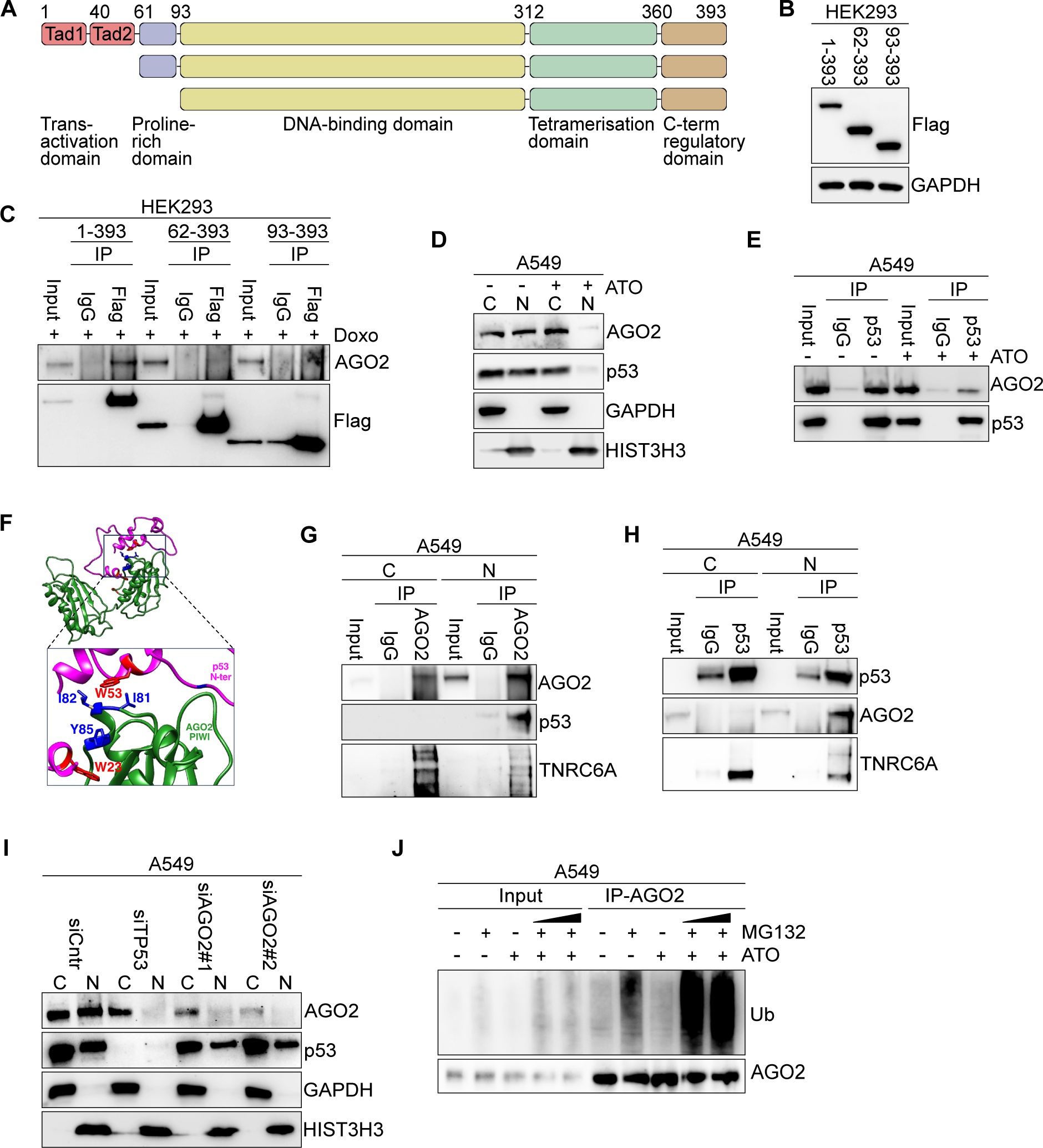
Tetrameric p53 protects nuclear AGO2 from proteasomal degradation. **(A)** Schematic diagram of human p53 protein and N-terminal truncated Flag-tagged p53 isoforms used in the study. **(B)** Representative Flag immunoblots from HEK293 cells transfected with Flag-WT-p53 or N-terminally Flag-tagged p53 mutants. GAPDH served as a loading control. n=3 **(C)** Flag immunoprecipitation (IP) from HEK293 cells transfected with plasmids expressing Flag-WT-p53 or N-terminally Flag-tagged p53 mutant. HEK293 cells were transfected with the p53 plasmids for 1 day and then treated with 1 μg/mL Doxorubicin (Doxo) for 1 more day. Representative immunoblots of AGO2 and Flag are indicated. n=3 **(D)** Representative AGO2 and p53 immunoblots from cytoplasmic (C) and nuclear (N) lysates in A549 cells treated with 0.5 μg/mL arsenic trioxide (ATO) for 24 hours. GAPDH served as a cytoplasmic marker and HIST3H3 served as nuclear marker. n=3 **(E)** p53 immunoprecipitation (IP) from A549 cells treated with 0.5 μg/mL Arsenite trioxide (ATO) for 24 hours. Representative immunoblots of AGO2 and p53. n=3 **(F)** Docking model of the PIWI domain of AGO2 with the N-terminal region of p53 was generated using the HDOCK server. The PIWI domain is depicted in green, and the N-terminal region of p53 is shown in pink (model 4) in ribbon representation. **(G)** AGO2 immunoprecipitation (IP) from cytoplasmic (C) and nuclear (N) lysates in A549 cells. Representative immunoblots of AGO2, p53 and TNRC6A. n=3 **(H)** p53 immunoprecipitation (IP) from cytoplasmic (C) and nuclear (N) lysates in A549 cells. Representative immunoblots of AGO2, p53 and TNRC6A. n=3 **(I)** Representative AGO2 and p53 immunoblots from cytoplasmic (C) and nuclear (N) lysates in A549 cells treated with siRNAs for 24 hours. siCntr: control scramble siRNAs; siTP53: siRNA specific for TP53; and siAGO2: two different siRNAs specific for AGO2. GAPDH served as a cytoplasmic marker and HIST3H3 served as nuclear marker. n=3 **(J)** AGO2 immunoprecipitation (IP) from A549 cells treated with 2 μg/ml MG132 for 2 hours before additional 24 hours of treatment with 0.5 μg/ml arsenic trioxide (ATO). Representative immunoblots of ubiquitin (Ub) and AGO2. n=3

Activated p53 tetramerizes in the nucleus ^39^ and therefore we next wanted to test if the tetrameric form is essential to interact with AGO2. In SK-N-BE(2) cells, known to possess intact *TP53* transcriptional activity, AGO2:p53 interacts in the nucleus (**Sup Fig 2C**) ^40^. To explore the effect of p53 monomerization, we treated SK-N-BE(2) cells with increasing concentrations of arsenic trioxide (ATO), a known inducer of p53 monomerization, for 24 hours ^41^. Remarkably, we observed a significant reduction in nuclear p53 localization with 0.1 μg/ml ATO treatment, and complete displacement of nuclear p53 at 0.5 μg/ml (**Sup Fig 3B**). Strikingly, AGO2 was also displaced from the nucleus upon treatment with 0.5 μg/ml ATO (**Sup Fig 3B**). Similar results were observed in A549 lung carcinoma cells (**Fig 3D**), HEK293T (**Sup Fig 3C**) and MCF7 breast cancer cells (**Sup Fig 3D**), where nuclear AGO2 and p53 accumulate in the cytoplasm after ATO treatment. Being able to manipulate p53 into its monomeric state, we next examined whether AGO2:p53 can interact under these conditions. Consequently, we performed co-immunoprecipitation assays in the above-mentioned cell lines, using whole cell lysates, in the presence or absence of ATO, and found that ATO-mediated monomerization of p53 reduces its capability to interact with AGO2 (**Fig 3E, Sup Fig 3E,F**), highly suggestive of the fact that p53 and AGO2 only interact in the nucleus.

Since the PIWI domain of AGO2 interacts with both p53 and TNRC6 at tryptophan-rich regions ^37^, we next investigated if these interactions are mutually exclusive. Therefore we docked the PIWI domain with the T6B region of TNRC6B and N-terminal region of p53 using the HDOCK server ^42^. The docking results indicated that AGO2 can bind to the N-terminal region of p53 and the T6B region of TNRC6 at different positions on the PIWI domain, depending on structural conformation of the proteins (**Fig 3F**, **Sup Table 2**). This suggests that the proteins may bind independently. To experimentally validate the computational models, we performed AGO2 or p53 immunoprecipitation assay from cytoplasmic and nuclear fractions of A549 cells. Our data suggest that AGO2 interacts with both p53 and TNRC6 in the nuclear fraction (**Fig 3G,H**) but, as expected, exclusivly with TNRC6 in the cytoplasm (**Fig 3G,H**).

Having found a robust interaction between AGO2:p53, we next wanted to gauge for its biological significance. To test whether p53 plays a role in the stability of AGO2, we silenced *TP53* using siRNAs. *TP53* silencing resulted in a decrease in AGO2 protein levels in A549 (**Fig 3I**), MCF7 (**Sup Fig 3G**) and HEK293T cells (**Sup Fig 3H**), suggesting that p53 may indeed stabilize AGO2 in the nucleus. However, silencing AGO2 in the same cells did not lead to reciprocal instability of p53 (**Fig 3I, Sup Fig 3G,H**) further corroborating the specificity of p53 mediated stability of AGO2. To confirm these findings, we employed p53 low-expressing MCF7 cells (hereafter referred to as TP53L cells) ^43^. Biochemical fractionation assays in wildtype (WT) and TP53L MCF7 cells revealed a similar downregulation of AGO2 protein levels, specifically in the nucleus, upon loss of p53 (**Sup Fig 3I**).

So far, we demonstrated that tetrameric p53 is needed for an interaction with AGO2 in the nucleus, however, it remained unclear whether p53 is needed for nuclear import of AGO2 or to stabilize AGO2 within the nucleus. Therefore, we tested if AGO2 is degraded by the ubiquitin-proteasome system when p53 is monomeric, by combining ATO treatment with the proteasome inhibitor MG132. In cells treated with MG132 and ATO, we found that AGO2 co-immunoprecipitated with anti-ubiquitin antibodies, indicating that AGO2 is tagged with ubiquitin for proteasome degradation in the absence of nuclear p53 (**Fig 3J, Sup Fig 3J**). Having previously observed that AGO1 and AGO3 also translocated into the nucleus in IAV-infected cells, we performed STRING analysis and identified that p53 may form complex with AGO1 and AGO2, but not with AGO3 and AGO4 (**Sup Fig 3K**). Therefore, we also evaluated if AGO1 is susceptible to p53 mediated degradation. Our results revealed that AGO1 co-immunoprecipitated with anti-ubiquitin antibodies in cells treated with MG132 and ATO, indicating that also AGO1 is degraded in the absence of nuclear p53 (**Sup Fig 3L**).

Collectively, our findings demonstrate an interaction between p53 and AGO2 in the nucleus, mediated by tetrameric p53. The interaction with p53 stabilizes AGO2 and protects it from proteasomal degradation.

### Nuclear AGO2 facilitates viral infection

Having observed a significant influx of both AGO2 and p53 into the nucleus upon viral infection and that p53 stabilizes AGO2 in the nucleus, we next aimed at determining if and how this phenomenon would influence viral infection. Specifically, we addressed whether nuclear AGO2:p53 serves as a proviral or an antiviral mechanism. To confirm whether the lack of p53 inhibits AGO2 translocation to the nucleus upon viral infection, we generated *TP53* knockout (KO) HEK293 cells by CRISPR-CAS9 (**Sup Fig 4A**). HEK293 cells infected with PR8-IAV at MOI 2 or MOI 10 translocated AGO2 into the nucleus (**Fig 1C**, **Fig 4A**), but the nuclear localization of AGO2 was lost in infected TP53 KO HEK293 cells (**Fig 4A**, **Sup Fig 4B**). Similarly, in WT MCF7 cells AGO2 predominantly localized in the nucleus, regardless of viral infection dose (**Sup Fig 4C**). In contrast, AGO2 remained cytoplasmic in TP53L MCF7 cells, irrespective of viral infection load (**Sup Fig 4D**). Furthermore, we measured viral gene expression in the above mentioned cells and observed a significant reduction of viral mRNA levels in TP53 KO HEK293 ad TP53L MCF7 cells (**Fig 4B, Sup Fig 4E**). To further verify that RNA expression of viral genes was correlated with viral infectivity, we infected WT and TP53 KO HEK293 cells, as well as WT and TP53L MCF7 cells, with PR8-mCherry virus and measured mCherry expression in infected cells using flow cytometry. TP53 KO HEK293 cells exhibited reduced viral replication (**Fig 4C**), which was was also evident in TP53L MCF7 cells (**Sup Fig 4F**). Overall, the reduced mRNA expression of NS1 and HA observed upon viral infection correlated with the lack of AGO2:p53 nuclear translocation (**Fig 4A, Sup Fig 4B-D**). Finally, we investigated the reversibility of this phenomenon by overexpressing p53 in TP53 KO HEK293 and in TP53L MCF7 cells. p53 overexpression partially restored viral gene expression (**Fig 4D, Sup Fig 4G**) and also promoted AGO2 nuclear accumulation (**Fig 4E, Sup Fig 4H**). Our data indicated that IAV infection is facilitated through the nuclear localization of AGO2:p53 complexes, however, it could not distinguish between a key role of either nuclear AGO2 or p53 in mediating the proviral outcome. Therefore, to test if AGO2 is essential in facilitating viral infectivity, we silenced *AGO2* in HEK293 cells by siRNAs, infected the cells with IAV and finally measured the mRNA levels of viral genes by qRT-PCR. It is important to consider that silencing *AGO2* did not affect the stability of nuclear p53 (**Fig 3I, Sup Fig 3G,H**). We found that mRNA levels of viral genes decreased when AGO2 was silenced in WT HEK293 cells (**Fig 4F, Sup Fig 4I-K).** Our results were recapitulated in A549 cells where we silenced either *AGO2* or *TP53* by siRNAs upon IAV infection and found that viral mRNA was significantly decreased (**Fig 4G**). Interestingly, silencing *AGO2* in TP53 KO HEK293 cells, infected with PR8-IAV, did not affect the levels of HA or NS1 viral mRNAs given the inability of AGO2 to translocate to the nucleus without p53 (**Sup Fig 4L**). Lastly, to confirm the role of NS1, we infected HEK293 cells with either PR8 or PR8-NS1_1-124_ mutant and silenced either p53 or AGO2. While silencing either p53 or AGO2 resulted in reduced viral mRNAs upon PR8 infection, it did not influence infectivity after PR8-NS1_1-124_ mutant infection, thus supporting the crucial role of NS1 (**Fig 4H,I**).

**Figure 4:**
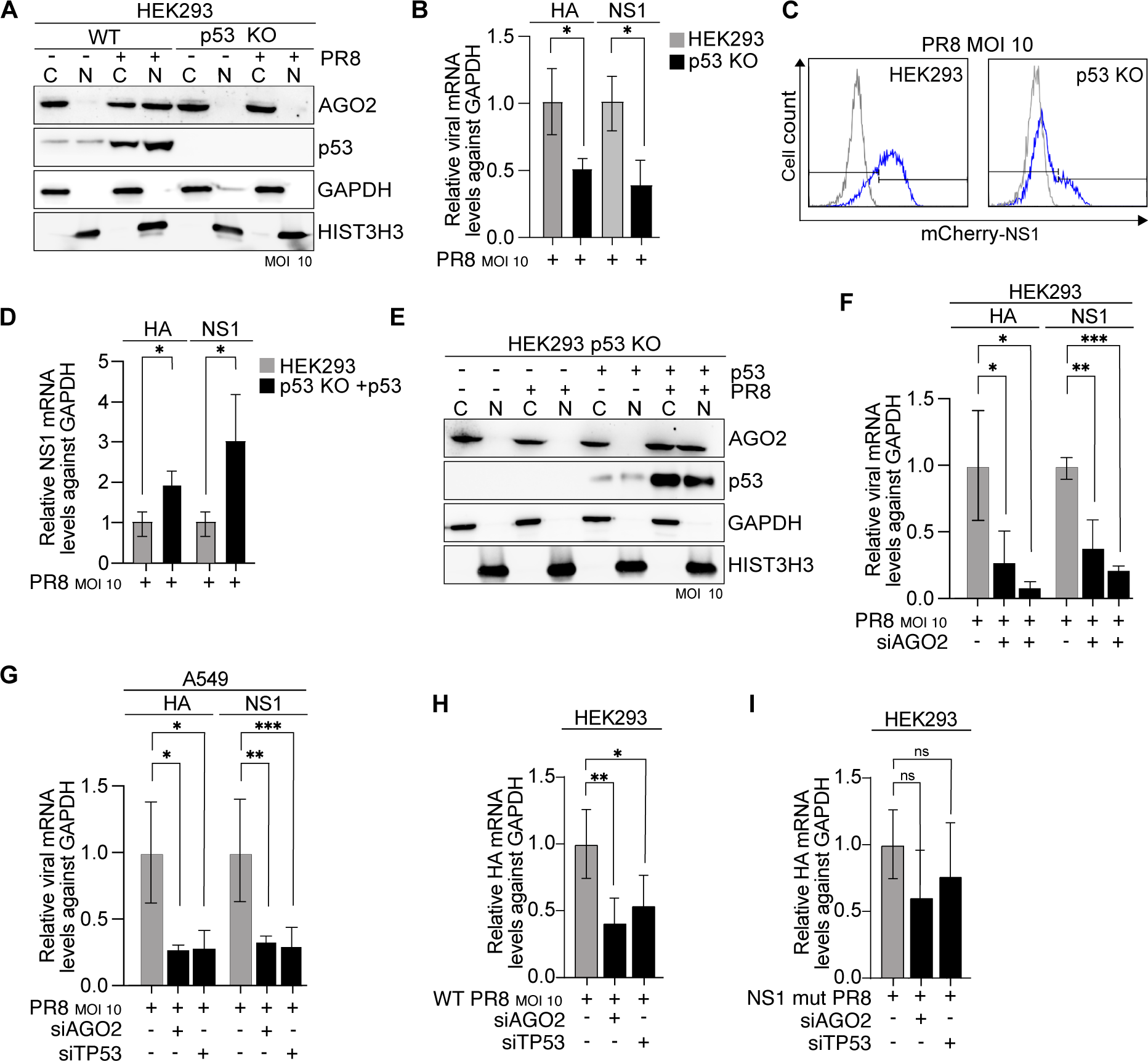
Nuclear AGO2 supports viral replication. **(A)** Representative AGO2 and p53 immunoblots from cytoplasmic (C) and nuclear (N) lysates in WT and TP53 KO HEK293 cells infected with PR8 virus at MOI 10 for 16 hours. GAPDH served as a cytoplasmic marker and HIST3H3 served as nuclear marker. n=3 **(B)** Relative expression, as measured by RT-qPCR, of HA and NS1 mRNA levels in WT and TP53 KO HEK293 cells upon infection with PR8 virus at MOI 10 for 16 hours. GAPDH was used as a reference gene. Bars are mean and error bars represent ± SD. ^∗^ p<0.05 by unpaired t-test. n=3 **(C)** Flow cytometry analysis of NS1-mCherry protein expression in WT and TP53 KO HEK293 cells upon infection with PR8 virus at MOI 10 for 16 hours. White histogram shows mock-infected cells while blue histogram is PR8-infected. n=3 **(D)** Relative expression, as measured by RT-qPCR, of HA and NS1 mRNA levels in WT and TP53 KO HEK293 cells. TP53 KO HEK293 cells were transfected with Flag-WT-p53 expressing plasmids for 24 hours. Subsequently, both TP53 KO HEK293 and TP53 KO HEK293 cells overexpressing WT p53 transiently were infected with PR8 virus at MOI 10 for 16 additional hours. GAPDH was used as a reference gene. Bars are mean and error bars represent ± SD. ^∗^ p<0.05 by unpaired t-test. n=3 **(E)** Representative AGO2 and p53 immunoblots from cytoplasmic (C) and nuclear (N) lysates in TP53 KO HEK293 cells transfected with Flag-WT-p53 expressing plasmids for 24 hours and infected with PR8 virus at MOI 10 for 16 additional hours. GAPDH served as a cytoplasmic marker and HIST3H3 served as nuclear marker. n=3 **(F)** Relative expression, as measured by RT-qPCR, of NS1 and HA mRNA levels in HEK293 cells treated with two different siRNAs against AGO2 (siAGO2) for 48 hours. 16 hours before the end of incubation, cells were infected with PR8 virus at MOI 10. GAPDH was used as a reference gene. Bars are mean and error bars represent ± SD. ^∗^ p<0.05, ^∗∗^ p<0.01, ^∗∗∗^ p<0.001 by unpaired t-test. n=3 **(G)** Relative expression, as measured by RT-qPCR, of NS1 and HA mRNA levels in A549 cells treated with siRNAs against AGO2 (siAGO2) or TP53 (siTP53) for 48 hours. 16 hours before the end of incubation, cells were infected with PR8 virus at MOI 10. GAPDH was used as a reference gene. Bars are mean and error bars represent ± SD. ^∗^ p<0.05, ^∗∗^ p<0.01, ^∗∗∗^ p<0.001 by unpaired t-test. n=3 **(H)** Relative expression, as measured by RT-qPCR, of HA mRNA levels in HEK293 cells treated with siRNAs against AGO2 (siAGO2) or TP53 (siTP53) for 48 hours. 16 hours before the end of incubation, cells were infected with WT PR8 virus at MOI 10. GAPDH was used as a reference gene. Bars are mean and error bars represent ± SD. ^∗^ p<0.05, ^∗∗^ p<0.01 by unpaired t-test. n=3 **(I)** Same as in **(H)** except cells were infected with PR8 virus expressing mutant NS1 at MOI 10 were. n=3

To summarize, we showed that mRNA levels of viral genes was similarly reduced after TP53 KO and siAGO2 (which had nuclear p53) in IAV-infected HEK293 cells, thus confirming an essential role for nuclear AGO2 but not p53 in the increased viral gene expression. Overall, we discovered a clear link between nuclear AGO2 localization and viral mRNA levels, indicating a proviral role of nuclear AGO2.

### Nuclear AGO2 downregulates antiviral type-I interferon response

Upon IAV infection, innate immunity is the first line of host defense, and the immediate immune response is mediated by type-I IFN. Type-I IFN are rapidly produced by infected cells to trigger an antiviral state, thus inhibiting viral replication ^44^. Given that AGO2:p53 nuclear translocation increased viral titer, we therefore hypothesized that AGO2:p53 nuclear translocation might have a proviral function by downregulating innate immune responses. First, we assessed the steady state levels of IFN-Is by measuring *IFNB* mRNA levels ^45^ by qRT-PCR in HEK293 vs HEK293T cells. Interestingly, we observed significantly higher levels of *IFNB* in HEK293 cells (nuclear AGO2 negative) compared to HEK293T cells (nuclear AGO2 positive) (**Fig 5A**), indicative of distinct IFN-I regulation between the two cell lines.

**Figure 5:**
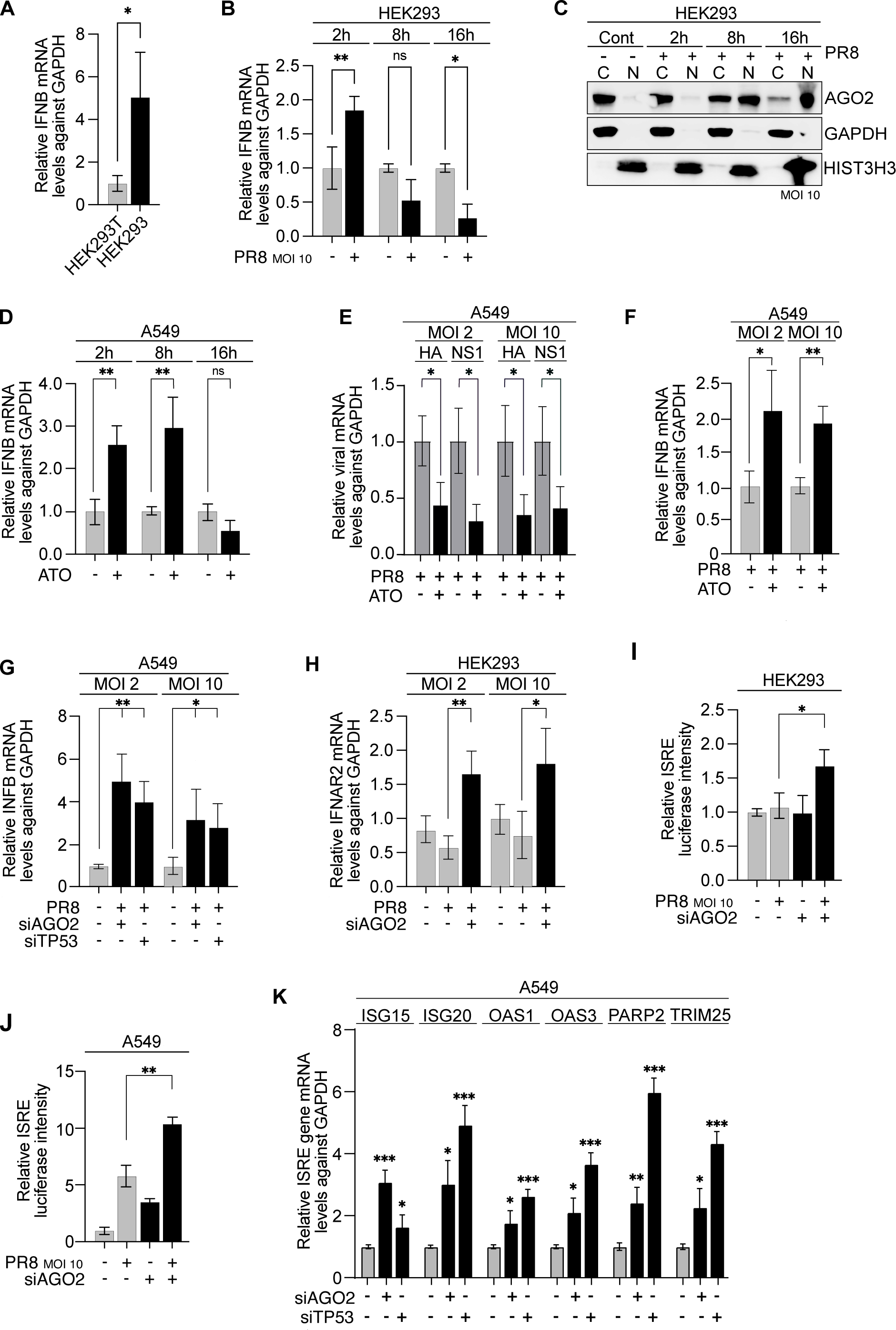
Nuclear AGO2 downregulates *IFNB* and other type-I-IFN related genes. **(A)** Relative expression, as measured by RT-qPCR, of IFNB mRNA levels in HEK293T and HEK293 cells. GAPDH was used as a reference gene. Bars are mean and error bars represent ± SD. ^∗^ p<0.1 by unpaired t-test. n=3 **(B)** Relative expression, as measured by RT-qPCR, of IFNB mRNA levels in HEK293 cells infected with PR8 virus at MOI 10 for 2, 8 or 16 hours. GAPDH was used as a reference gene. Bars are mean and error bars represent ± SD. ^∗^ p<0.05, ^∗∗^ p<0.01 by unpaired t-test. n=3 **(C)** Representative AGO2 immunoblots from cytoplasmic (C) and nuclear (N) lysates in HEK293 cells infected with PR8 virus at MOI 10 for 2, 8 or 16 hours. GAPDH served as a cytoplasmic marker and HIST3H3 served as nuclear marker. n=3 **(D)** Relative expression, as measured by RT-qPCR, of IFNB mRNA levels in A549 cells treated with 0.5 μg/mL arsenic trioxide (ATO) for 2, 8 or 16 hours. GAPDH was used as a reference gene. Bars are mean and error bars represent ± SD. ^∗∗^ p<0.01 by unpaired t-test. n=3 **(E)** Relative expression, as measured by RT-qPCR, of HA and NS1 mRNA levels in A549 cells treated for 2 hours with 0.5 μg/ml arsenic trioxide (ATO)) or vehicle and infected with PR8 virus at MOI 2 or MOI 10 for 16 hours. GAPDH was used as a reference gene. Bars are mean and error bars represent ± SD. ^∗^ p<0.05 by unpaired t-test. n=3 **(F)** same as in **(E)** but IFNB mRNA level were measured by RT-qPCR. n=3 **(G)** Relative expression, as measured by RT-qPCR, of IFNB mRNA levels in A549 cells treated with siRNAs against TP53 or AGO2 for 48 hours. 16 hours before the end of incubation, cells were infected with PR8 virus at MOI 2 or MOI 10. GAPDH was used as a reference gene. Bars are mean and error bars represent ± SD. ^∗∗∗^ p<0.001, ^∗∗∗∗^ p<0.0001 by unpaired t-test. n=3 **(H)** Relative expression, as measured by RT-qPCR, of IFNAR2 mRNA levels in HEK293 cells treated with siRNA against AGO2 for 48 hours. 16 hours before the end of incubation, cells were infected with PR8 virus at MOI 2 or MOI 10. GAPDH was used as a reference gene. Bars are mean and error bars represent ± SD. ^∗^ p<0.05 by unpaired t-test. n=3 **(I)** Normalized lucifersase signal of ISRE-transfected HEK293 cells, treated with siRNA against AGO2 for 48 hours. 16 hours before the end of incubation, cells were infected with PR8 virus at MOI 10. GAPDH was used as a reference gene. Bars are mean and error bars represent ± SD. ^∗^ p<0.05 by unpaired t-test. n=3 **(J)** same as in **(I)** but in A549 cells. n=3 **(K)** Relative expression, as measured by RT-qPCR, of ISG15, ISG20, OAS1, OAS3, PARRP12, and TRIM25 mRNA levels A549 cells treated with siRNA against AGO2 for 48 hours. 16 hours before the end of incubation, cells were infected with PR8 virus at MOI 10. GAPDH was used as a reference gene. Bars are mean and error bars represent ± SD. ^∗^ p<0.05 by unpaired t-test. n=3

Subsequently, we examined the dynamics of *IFNB* levels in HEK293 cells at three distinct time points following PR8 infection. We observed an initial increase in *IFNB* at 2 hours post-infection, followed by a decrease at 8- and 16-hours post-infection, where the levels were lower than baseline (**Fig 5B**). In the same time frame mentioned above, we also performed biochemical fractionation assays. Strikingly, the decline in *IFNB* production coincided with the translocation of AGO2 into the nucleus at 8 hours post-infection, suggesting a potential contribution of nuclear AGO2 in the downregulation of antiviral IFN-I production (**Fig 5C**).

To explore whether the regulation of IFN-I expression is a general feature in cells positive for nuclear AGO2, we measured *IFNB* in WT and TP53L MCF7 cells. Notably, we observed low levels of *IFNB* expression in WT MCF7 cells (with nuclear AGO2), while the TP53L MCF7 cells, lacking nuclear AGO2, displayed significantly higher *IFNB* levels (**Sup Fig 5A**). Having observed that the lack of p53, and consequent lack of nuclear AGO2, allows for the expression of *IFNB*, we next reasoned that we should be able to recapitulate the above observation in nuclear AGO2 positive cells when p53 is monomeric and AGO2:p53 complex is excluded from the nucleus (**Fig 3D, Sup Fig 3C,D**). To investigate the impact of p53 monomerization on IFN-I expression we measured *IFNB* levels in A549 and HEK293T cells treated with ATO and observed a significant increase in *IFNB* (**Fig 5D, Sup Fig 5B**), which is likely due to the exclusion of AGO2:p53 from the nucleus. Furthermore, as expected, A549, HEK293T and MCF7 cells treated with ATO and infected with IAV exhibited reduced viral gene expression (**Fig 5E, Sup Fig 5C,D**), linked with rescued *IFNB* levels (**Fig 5F, Sup Fig 5E,F**). In addition, to further elucidate the role of AGO2:p53 axis in regulating *IFNB* expression, we performed siRNA-mediated knockdown of *AGO2* and *TP53* in A549 and HEK293T cells and measured *IFNB* mRNA levels (**Fig 5G, Sup Fig 5G**). Silencing either *AGO2* or *TP53* led to an increase in *IFNB* mRNA levels (**Fig 5G, Sup Fig 5G**). Importantly, p53 nuclear localization is not compromised with AGO2 silencing (**Fig 3I, Sup Fig 3G,H**) hence suggesting a direct role of nuclear AGO2 in *IFNB* regulation.

Type-I IFNs, which are produced by infected cells, trigger a signaling cascade that timely leads to an antiviral state by promoting the expression of interferon simulating response elements (ISRE), via IFNAR1 and IFNAR2 stimulation ^44^. Therefore, we next wanted to gauge whether increase in *IFNB* triggered downstream activation of IFNAR-mediated ISRE pathway after IAV infection. Importantly, silencing of *AGO2* by siRNA resulted in a significant increase of *IFNAR2* expression in HEK293 cells infected with IAV (**Fig 5H**). To assess the downstream effects of interferon response, we measured the expression of ISRE. Utilizing a luciferase reporter assay in both IAV-infected and control HEK293 cells, we found that the ISRE expression was significantly increased in IAV-infected cells only when AGO2 was silenced (**Fig 5I**). ISRE measurements in A549 (nuclear AGO2 positive) demonstrated an increase of ISRE in both infected and uninfected cells, upon AGO2 silencing (**Fig 5J**). Luciferase results were confirmed by qPCR that showed the upregulation of several ISRE genes in IAV-infected cells upon AGO2 and TP53 silencing in both A549 and HEK293T cells (**Fig 5K, Sup Fig 5H**). Taken together, our findings support the notion that nuclear localization of AGO2 acts as a mechanism to suppress the induction of antiviral IFN-I and its signaling cascade, thereby facilitating viral infection. This resistance mechanism utilized by viruses highlights the complex interplay between viral pathogens and the host immune response.

### Type-I-IFN-pathway-related genes and *TRIM71* are negatively regulated by nuclear RNAi

As nuclear AGO2 translocation correlated with diminished interferon expression and enhanced viral gene expression, we hypothesized that nuclear RNAi may directly suppress type-I IFNs. First, to comprehensively understand the transcriptional dynamics of host cellular responses during IAV infection, we conducted RNA sequencing experiments in HEK293 cells post-IAV infection. The results of principal component analysis (PCA) demonstrated that IAV infection elicited a distinctive transcriptomic profile in HEK293 cells (**Sup Fig 6A**). Further, we identified 1773 differentially upregulated and 352 differentially downregulated RNA transcripts in response to IAV infection (**Sup Fig 6B** and **Sup Table 3**). As expected, gene ontology (GO) analysis of biological processes revealed that PR8 infection significantly induced defense response and regulation of immune effector processes (**Sup Fig 6C)**. Upregulated genes, associated with inflammation and innate immunity, included *ARRD3, ITGB1BP2, SOCS1, SOCS3, TRIM72, GADD45B,* and *CD68* (**Sup Fig 6B**). Notably, among the upregulated genes, *TRIM72*, and *SOCS1* emerged as potential inhibitors of the IFN-I response, with reported interactions with RIG-1, IFNB, and IFNAR, respectively ^46–48^. While *IFNB* itself could not be detected in the RNAseq dataset, many genes upstream of *IFNB*, including *MAVS*, *IRF3*, *TRIF3*, and *TRIF6,* were downregulated.

To explore the role of nuclear RNAi and define AGO targets at nucleotide resolution, we employed fPAR-CLIP assay ^49^ in two replicates in HEK293 cells with or without IAV infection, from either cytoplasmic or nuclear fractions and using the T6B peptide, which recognizes all four AGOs, to isolate AGO-bound RNAs ^50^ (**Sup Fig 6D**). Overall, we identified 41 743 cytoplasmic and 12 119 nuclear AGO1-4 binding sites in control cells (**Fig 6A**). At 16 hours post IAV infection, we identified 30 083 cytoplasmic and 78 665 nuclear AGO1-4 binding sites, a remarkable 6.5 fold increase of nuclear AGO-targets (**Fig 6A**). A clear difference in control *vs* IAV-infected nuclear AGO target occupancy was also evident by PCA analysis (**Sup Fig 6E**). Finally, when mapping the distribution of fPAR-CLIP sequence reads across target RNAs little changes were observed in AGO target occupancy obtained from the cytoplasmic fraction, with or without viral infection (**Fig 6A**). Contrary, the striking increase in nuclear AGO transcript clusters, suggested that, once AGOs enter the nucleus, they expand their binding preferences and interact with the pre-mRNA sequence. The predominant target occupancy was on intronic sites, but also at 3’UTR and CDS regions within the nucleus (**Fig 6A**).

**Figure 6:**
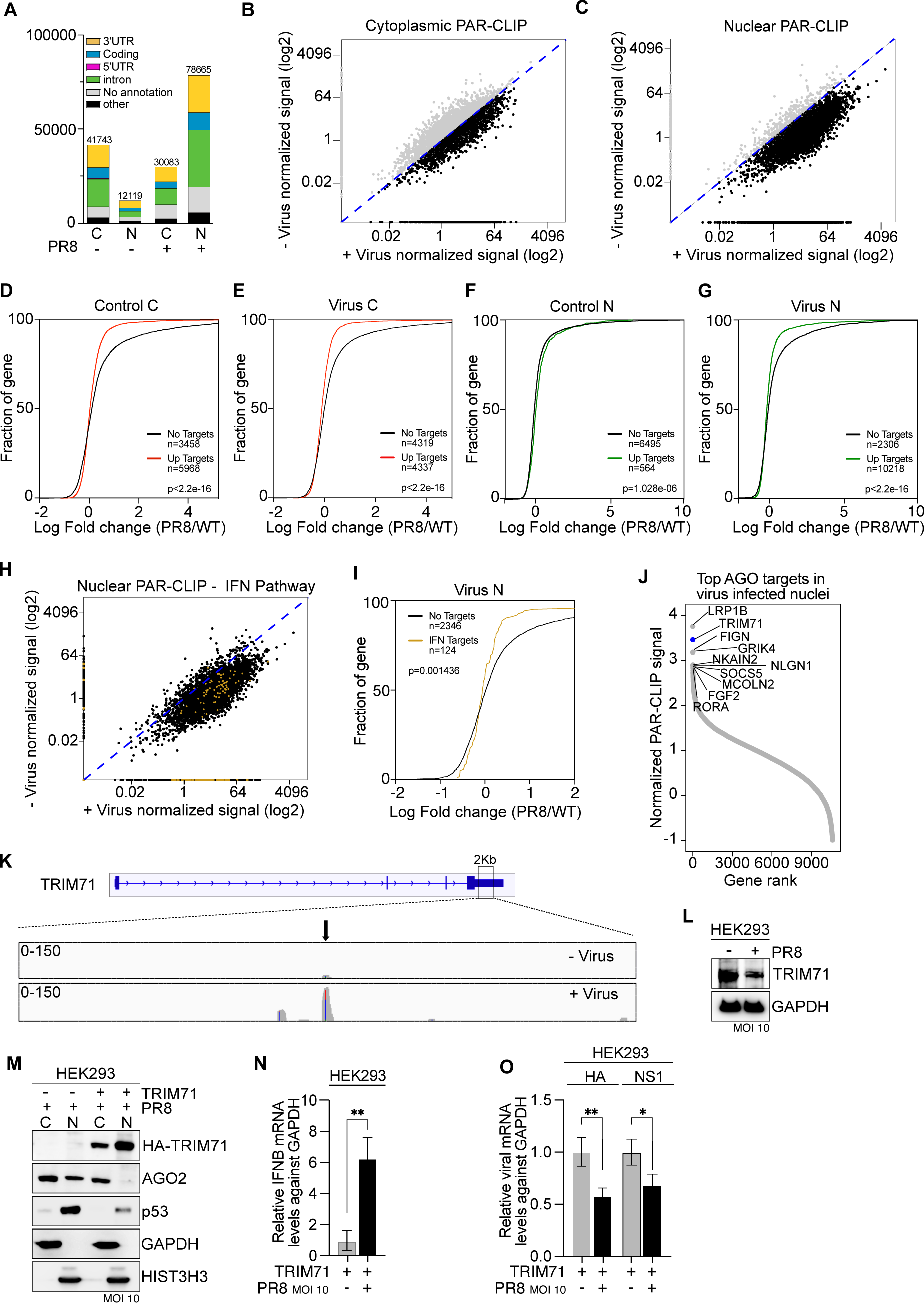
Nuclear AGO2 downregulates type-I IFN pathway genes and TRIM71 in IAV-infected cells. **(A)** Distribution of fPAR-CLIP sequence reads in clusters across target RNA across 3′UTR, coding sequence, 5′UTR, and introns from cytoplasmic (C) and nuclear (N) fractions of HEK293 cells infected with PR8 virus at MOI 10 for 16 hours. The graph shows the average of 2 independent experiments. **(B)** Scatter plot of cytoplasmic fPAR-CLIP log2 normalized signal in control (-Virus) and IAV infected (+Virus) HEK293 cells. Each dots represents a gene and is the average of 2 independent experiments. Blue dashed line shows a perfect correlation. Black dots are genes with higher PAR-CLIP signal in the IAV-infected sample while grey dots are genes with higher PAR-CLIP signal in the control sample. **(C)** same as in **(B)** but for nuclear fPAR-CLIP. **(D)** Cumulative distribution function of AGO1-4 targets in control cytoplasmic fraction compared to non-targets. Targets are the genes depicted as grey dots in (**B**) **(E)** Cumulative distribution function of AGO1-4 targets in virus infected cytoplasmic fraction compared to non-targets. Targets are the genes depicted as black dots in (**B**) **(F)** Cumulative distribution function of AGO1-4 targets in control nuclear fraction compared to non-targets. Targets are the genes depicted as grey dots in (**C**) **(G)** Cumulative distribution function of AGO1-4 targets in virus infected nuclear fraction compared to non-targets. Targets are the genes depicted as black dots in (**C**) **(H)** same graph as in **(C)** but with type-I IFN pathway genes highlighted in golden yellow. **(I)** same as in **(G)** but for type-I IFN pathway genes. **(J)** Rank plot showing AGO targets, ordered by normalized PAR-CLIP signal, in IAV infected, nuclear fraction of HEK293 cells. Each dot represents a gene and is the average of 2 independent experiments. **(K)** TRIM71 IGV track in nuclear fraction of in control (-Virus) and IAV infected (+Virus) HEK293 cells. **(L)** Representative TRIM71 immunoblots from whole cell lysates in HEK293 infected with PR8 virus at MOI 10 for 16 hours. GAPDH served as a loading control. n= 3 **(M)** Representative HA-TRIM71, AGO2 and p53 immunoblots from cytoplasmic (C) and nuclear (N) lysates in HEK293 cells transfected with HA-TRIM71 and infected with PR8 virus at MOI 10 for 16 hours. GAPDH served as a cytoplasmic marker and HIST3H3 served as nuclear marker. n= 3 **(N)** Relative expression, as measured by RT-qPCR, of IFNB mRNA levels in HEK293 cells transfected with HA-TRIM71 and infected with PR8 virus at MOI 10 for 16 hours. GAPDH was used as a reference gene. Bars are mean and error bars represent ± SD. ∗∗ p<0.01 by unpaired t-test. n= 3 **(O)** Relative expression, as measured by RT-qPCR, of HA and NS1 mRNA levels in HEK293 cells transfected with HA-TRIM71 and infected with PR8 virus at MOI 10 for 16 hours. GAPDH was used as a reference gene. Bars are mean and error bars represent ± SD. ∗ p<0.05, ∗∗ p<0.01 by unpaired t-test. n= 3

Next, we sought to investigate what targets the AGOs regulate upon viral infection. To better visualize shifts in AGO-binding within the different cellular fractions, we plotted the fPAR-CLIP signal, for all coding genes, in control (-virus) *vs* IAV-infected (+ virus) cells. While dots on the axis indicate unique binders, a shift from the dashed line (x=y) denotes enrichment in the number of AGOs binding to each transcript and/or the number of bound transcripts. The resulting scatter plots highlighted no changes in the cytoplasmic environment (**Fig 6B**) but a surge in AGO binding, within the nucleus, post-infection (**Fig 6C**). Genes identified as having different distribution in the scatter plots, between IAV infected and non infected samples, were defined as AGO2 targets. Taken together, the results demonstrate that, upon IAV infection, not only is there an enrichment in nuclear AGO targets, but that more AGOs are bound to each transcripts, upon viral infection.

Furthermore, to understand the effect of AGO-binding, we analyzed AGO binding targets by cumulative distribution analysis, in relation to RNAseq experiment (**Sup Fig 6B**). Here, a shift of the cumulative distribution function (CDF) curve to the left means that a higher proportion of AGO targets are effectively suppressed, indicating that the RNAi machinery is active and efficient. Consistent with the canonical function of cytoplasmic RNAi, we found that AGO1-4 suppressed its best binding targets in the cytoplasm equally, regardless of IAV infection (**Fig 6D,E**). On the contrary, in the nuclei of control cells, the differences in expression fold change between AGO targets and non-targets are minor, as shown in the CDFs and supported by the fact that AGOs are not present in the nucleus in steady state HEK293 cells (**Fig 6F**). However, we observed potent negative gene regulation of nuclear AGO fPAR-CLIP targets, after IAV infection (**Fig 6G**).

Our previous data (**Fig 5**) suggested that AGO2 nuclear translocation correlated with a decrease in type-I IFN pathway, which is crucial for antiviral responses. To test whether nuclear AGOs had a direct effect on genes specifically involved in type-I IFN response, we highlighted targets specifically involved in IFN-Is response ^51^ in the scatter plot and performed CDF using those specific targets (**Fig 6H,I**). Indeed, 124 out of 131 genes were nuclear AGO targets and were substantially downregulated by nuclear RNAi (**Fig 6H,I**).

To further gain insight into what genes AGO1-4 targets in the nucleus during viral infection, we ranked the fPAR-CLIP targets of the AGOs (**Fig 6J, Sup Table 4**). Notably, among the top AGO-bound transcripts in the nuclear fraction from viral-infected cells, three of the top ten targets were in the 3’UTR and intronic regions of *LRP1B*, *TRIM71*, and *SOCS5* (**Fig 6J**). These molecules are well known for their significant roles in the immune response ^11,52,53^. Of particular interest is *TRIM71* (also referred to as LIN41), renowned for its positive influence on the IFNβ and ISRE responses but also directly inducing AGO2 and p53 degradation processes ^11,54,55^. Further elucidation from a STRING analysis revealed that AGO2 predominantly interacts with TRIM family proteins that plays a positive role in type-I interferon pathway, especially with TRIM56, and TRIM71 (**Sup Fig 6F**). Our fPAR-CLIP data, interpreted via the IGV software ^56^, clearly showed an enhanced AGO2 binding to *TRIM71*, *LRP1B*, *SOCS5*, *IFNAR2* and *TRIM56* 3’UTR site following IAV infection, compared to controls (**Fig 6K, Sup Fig 6G-J**). Substantiating this observation, post IAV-infection, there was a significant decrease in TRIM71 and IFNAR2 protein levels (**Fig 6L, Sup Fig 6K**). Finally, further examination of RNA-seq data showed downregulation of *TRIM71* mRNA (log2FC −0.38), highlighting the dynamic response of host cellular machinery to viral infection.

Our experimental data indicated that IAV infection promotes AGO2 nuclear translocation, and the fPAR-CLIP results suggest two possible complimentary functions of nuclear RNAi. 1) Direct silencing of genes involved in the type-I IFN pathways and, 2) targeting of TRIM71 to block its direct effect on IFN response but also to prevent AGO2 and p53 degradation. To experimentally test the latter, we overexpressed TRIM71 transiently in IAV-infected HEK293; indeed, we observed a marked reduction in nuclear AGO2 levels (**Fig 6M**). This coincided with diminished p53 levels, aligning with the recognized role of TRIM71 as a p53 and AGO2 E3 ligase ^11,54,55^. Consequently, there was an increase in *IFNB* (**Fig 6N**) and other type-I IFN related genes mRNA levels (**Sup Fig 6L**), and a decrease in viral mRNA (**Fig 6O**), attributable to the innate immune function of TRIM71 and decreased presence of nuclear AGO2.

These insights suggest a nuanced viral strategy that involves AGO2-mediated gene silencing combined with TRIM71 and IFNAR2 targeting. The culmination of these actions dampens the type-I IFN pathway, allowing the virus to adeptly evade host immune defenses. In summary, our integrated approach combining RNA sequencing and fPAR-CLIP demonstrates that nuclear AGO2 is crucial for the virus to subvert host immune responses and to ensure a successful infection.

### AGO2 targets TRIM71 through Let-7 miRNAs, upon IAV infection

AGO proteins exert their gene regulation by miRNAs. The miRNAs guide AGO to its target RNA transcript and Watson-Crick basepairing allows for hybridization with the target, RISC assembly and recruitment of effector complex for gene regulation ^57^. Having determined that TRIM71 is one of the top targets of AGO2 in the nucleus upon viral infection, we next wanted to evalute which miRNAs are responsible for TRIM71 targeting. Using IGV software we identified that AGO2 binds to two specific regions on the TRIM71 3’UTR (**Fig 6K**), which are associated with the Let-7/98/4458/4500 and miR181abcd/4262 families (**Fig 7A**,). Given the established role of Let-7 in cellular senescence, which is also caused by viral infection, we focused our investigation on Let-7 ^58,59^. To study the regulatory mechanism of Let-7 on AGO2 and its impact on cellular phenotype, we utilized LIN28, which is known to inhibit the processing of Let-7 precursors, thus controlling the levels of mature Let-7 miRNAs ^60^. First, we overexpressed LIN28A and LIN28B in MCF7 cells and observed that LIN28A effectively prevents the nuclear accumulation of AGO2 (**Sup Fig 7A**). We further tested the effect of LIN28A overexpression on AGO2 localization in the context of PR8 infection in A549 and HEK293 cells demonstrating that LIN28A does indeed reduces nuclear AGO2 accumulation (**Fig 7B, Sup Fig 7B**). NS1 protein levels were also deacreased in viral infected and LIN28A overexpressing cells, futher supporting that blocking nuclear AGO2 via Let-7 may reduce viral infection (**Fig 7C**). To confirm a role for the miRNAs, we measured the levels of Let-7c, Let-7f and Let-7g mature miRNA by qPCR and observed increased expression of Let-7c/f/g in PR8-infected cells, in contast to control cells, which was promptly reduced by LIN28A overexpression (**Fig 7D-F**). Together, these results provide compelling evidence that modulating Let-7 levels, through LIN28A overexpression, impacts AGO2 dynamics and potentially the cellular response to IAV infection.

**Figure 7:**
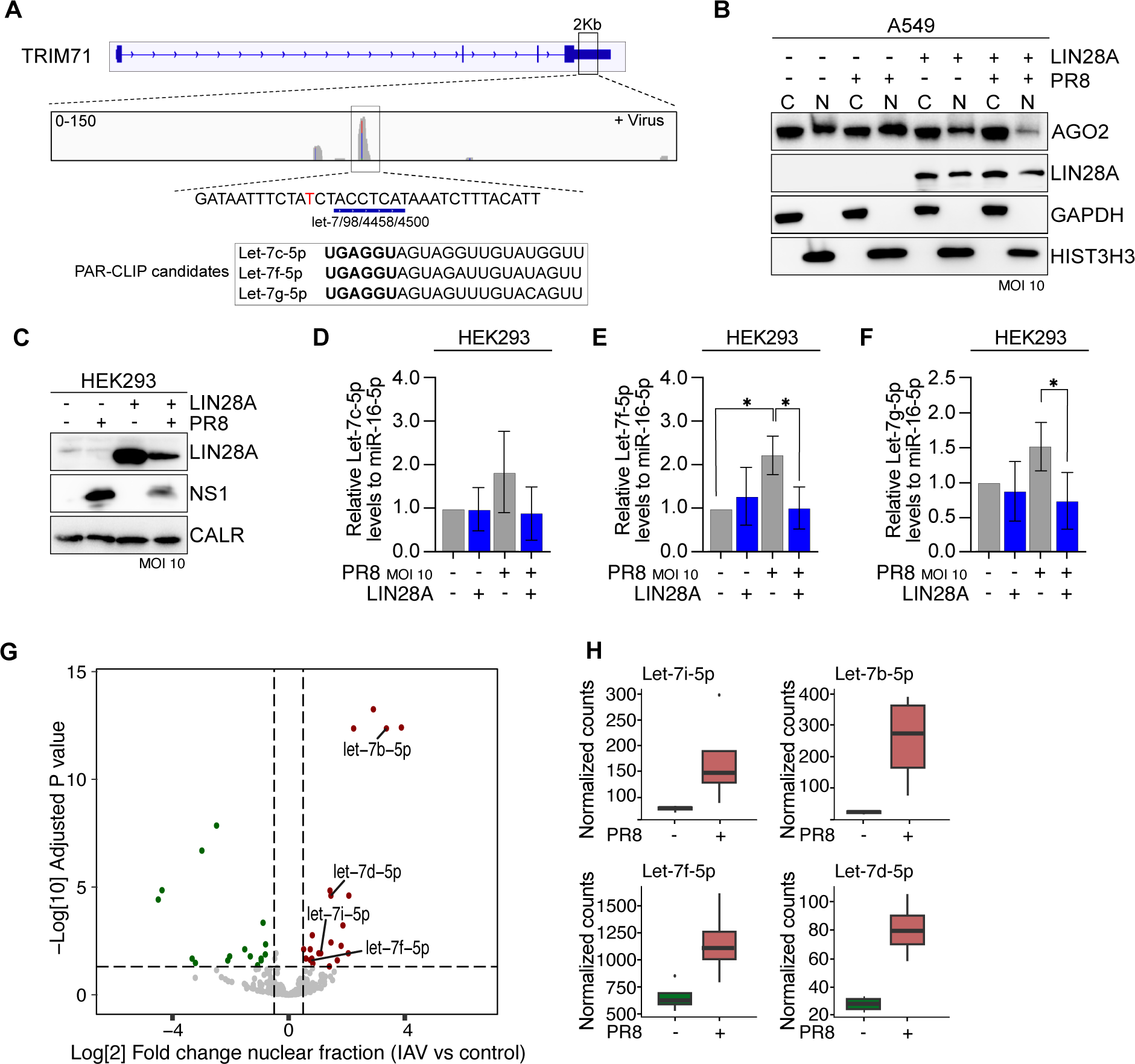
Let-7 miRNA, targeting TRIM71, are upregulated in the nucleus of IAV-infected cells. **(A)** TRIM71 IGV track in nuclear fraction of in IAV infected (+Virus) HEK293 cells. Indicated is the TRIM71 sequence, predicted Let-7 seed sequence and Let-7 PAR-CLIP targets. **(B)** Representative AGO2 and LIN28A immunoblots from cytoplasmic (C) and nuclear (N) lysates in A549 cells transfected with LIN28A expressing plasmid and infected with PR8 virus at MOI 10 for 16 hours. GAPDH served as a cytoplasmic marker and HIST3H3 served as nuclear marker. n=3 **(C)** Representative LIN28A and NS1 immunoblots from cytoplasmic (C) and nuclear (N) lysates in HEK293 cells transfected with LIN28A expressing plasmid and infected with PR8 virus at MOI 10 for 16 hours. CALR served as a loading control. n=3 **(D)** Relative expression, as measured by RT-qPCR, of Let-7c-5p RNA levels in HEK293 cells transfected with V5-LIN28A expressing plasmid and infected with PR8 virus at MOI 10 for 16 hours. miR-16-5p was used as a reference gene. Bars are mean and error bars represent ± SD. n= 3 **(E)** same as in **(D)** but for Let-7f-5p RNA. Bars are mean and error bars represent ± SD. ∗ p<0.05 by unpaired t-test. n= 3 **(F)** same as in **(D)** but for Let-7g-5p RNA. Bars are mean and error bars represent ± SD. ∗ p<0.05 by unpaired t-test. n= 3 **(G)** Volcano plot showing miRNAseq results of differentially expressed miRNAs from the nuclear fraction of PR8 infected and non infected HEK293 cells. In red are the upregulated genes while in green the downregulated. **(H)** Significantly differentially expressed Let7i/b/f/d-5p from the nuclear fraction of PR8 infected and non infected HEK293 cells.

To futher evaluate the link between nuclear AGO and gene silencing, we assessed the global miRNA profiles bound to AGO proteins in the presence or absence of viral infection. HEK293 cells were infected with PR8 virus, fractionated into cytoplasmic and nuclear fractions, and AGO-specific targets were enriched by AGO1-4 pulled down using the T6B peptide (**Sup Fig 7C**) before small RNA extraction and sequencing (**Sup Fig 7D**). Differential expression analysis demonstrated significant changes in the nuclear fraction upon infection, with significant upregulation of miRNAs, belonging to the Let-7 family, in the nuclear fraction (**Fig 7G,H, Sup Table 5**). Similar changes were not observed in the cytoplasmic fraction, thus confirming specific nuclear targeting of TRIM71 upon IAV infection (**Sup Fig 7E,F**). In order to exclude a direct effect of miRNA on viral genes, we mapped sequencing reads to the IAV genome and visualized targeting with IGV software. The tracks clearly showed no enrichment over background upon infection thus discounting a direct role of nuclear or cytoplasmic AGO2 in miRNA-mediated silencing of viral genes (**Sup Fig 7G**). Overall, we demonstrate a critical role for miRNAs of the Let-7 family, targeting TRIM71, in the AGO2-mediated nuclear regulation of IAV infectivity.

### Nuclear p53-AGO2 axis is involved in regulation of innate immunity and IAV infectivity *in vivo*

Our data demonstrates that AGO2 suppresses type-I IFN in the nucleus and that p53 is essential to mediate AGO2 nuclear translocation. To confirm these mechanisms *in vivo*, we intranasally (i.n.) administered IAV-PR8 to wild-type C57BL/6 mice. Lung tissues were harvested on days 1 and 3 post-infection (d.p.i.) (**Fig 8A**). As expected, mice infected with IAV exhibited markedly elevated viral mRNA levels in lung tissues when compared to controls at 3 d.p.i. (**Fig 8B, Sup Fig 8A**). Next, we isolated single cells from both control and IAV-infected lung tissues for biochemical fractionations assays. Healthy lung cells predominantly showed cytoplasmic AGO2 distribution. In contrast, a notable nuclear accumulation of AGO2 was observed in response to IAV infection (**Fig 8C**). To exclude that the positive AGO2 signal we observed in the nuclear fraction was due to the influx of immune cells in the infected lungs ^61^, we carried out a negative selection of CD45^+^ immune cells. We found that AGO2 translocated to the nucleus in both CD45^+^ and CD45^-^ cells, thus confirming that IAV infection induced AGO2 translocation in the epithelial and endothelial cells (**Sup Fig 8B**) and providing *in vivo* evidence supporting IAV-mediated AGO2 nuclear translocation. To strengthen the role of NS1 *in vivo*, we infected mice with PR8-NS1_1-124_ mutant and collected lungs at 1 and 3 d.p.i. (**Sup Fig 8C**). Mutant NS1 IAV infection did not induce AGO2 nuclear translocation (**Sup Fig 8D**) and, accordingly, mice exhibited lower viral titers (**Sup Fig 8E,F**) and heightened *Ifnb* levels (**Sup Fig 8G**).

**Figure 8:**
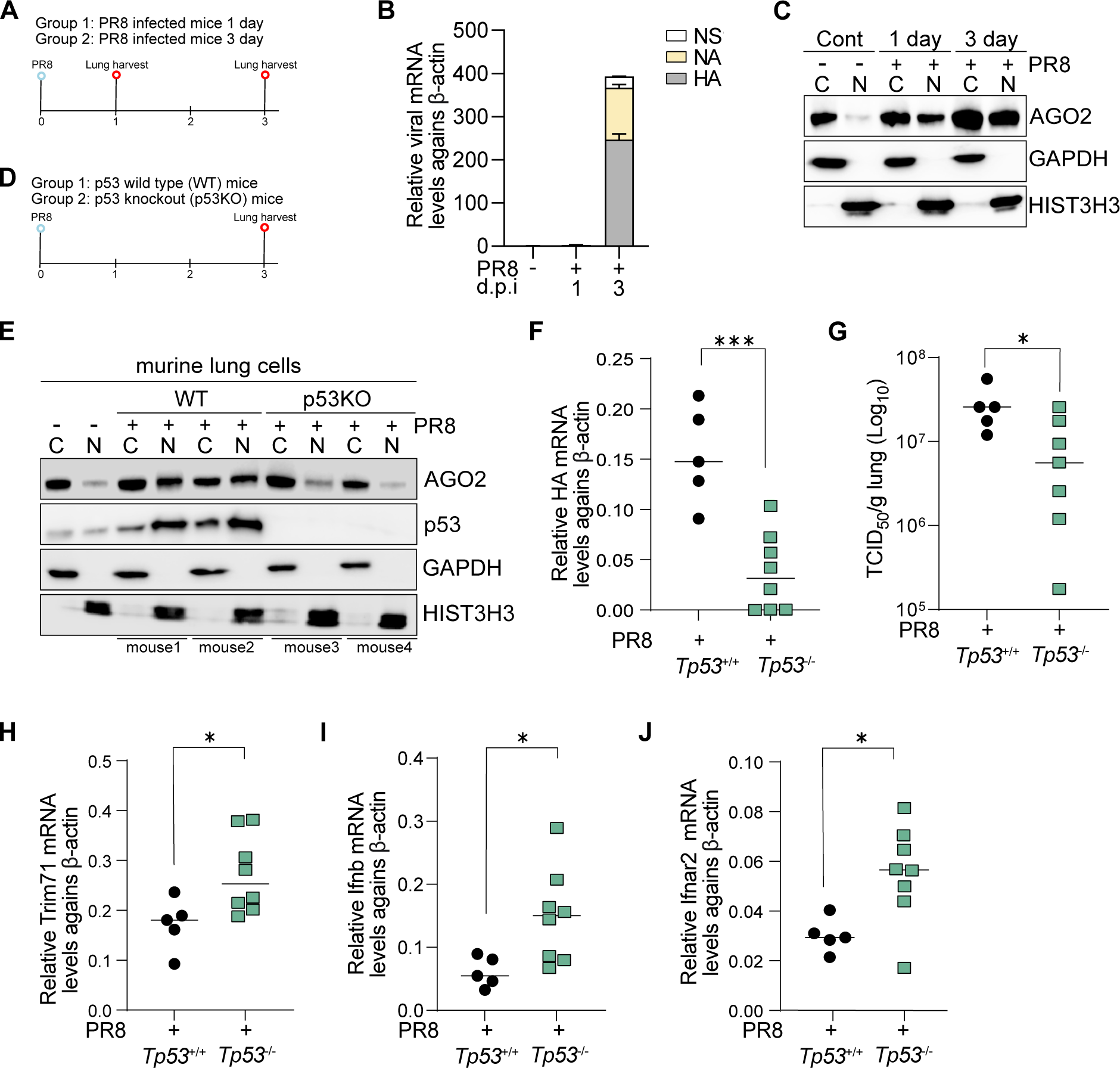
p53/AGO2 axis contributes to the decrease in IFN-related genes and increased viral titers *in vivo*. **(A)** Schematic representation of the experimental setup for experiments in **(B)** and **(C)**. Mice were infected i.n. with 2000 TCID_50_ PR8 at day 0 and lungs harvested at 1 and 3 days post-infection. **(B)** Relative expression, as measured by RT-qPCR, of NS1 and HA mRNA levels in lung cells isolated from mice at 1 or 3 days post-infection. β-actin was used as a reference gene. Error bars represent ± SD. **(C)** Representative AGO2 immunoblots from cytoplasmic (C) and nuclear (N) lysates in lung cells isolated from mice at 1 or 3 days post-infection. GAPDH served as a cytoplasmic marker and HIST3H3 served as nuclear marker. n= 2 independent experiments with 2 mice each **(D)** Schematic representation of the experimental setup for experiments in **(E-J)**. WT and *Tp53-/-* mice were infected i.n. with 2000 TCID_50_ PR8 and lungs harvested at 3 days post-infection. **(E)** Representative Ago2 and p53 immunoblots from cytoplasmic (C) and nuclear (N) lysates in lung cells isolated from WT and *Tp53^-/-^* mice infected with PR8. GAPDH served as a cytoplasmic marker and HIST3H3 served as nuclear marker. n=1 independent experiments with a total of 5 WT and 8 *Tp53^-/-^* mice. **(F)** Graph representing HA mRNA levels in lung cells isolated from WT and *Tp53^-/-^* mice infected with PR8. Shown are the individual mice with bar respresenting the mean and error bars represent ± SD. ^∗∗∗^ p<0.001 by unpaired t-test. n=1 independent experiments with a total of 5 WT and 8 *Tp53^-/-^*mice. **(G)** Graph representing log_10_ TCID_50_/g lung in WT and *Tp53^-/-^* mice with PR8 infection. Shown are the individual mice with bar respresenting the mean and error bars represent ± SD. ^∗^ p<0.05 by unpaired t-test. n=1 independent experiments with a total of 5 WT and 8 *Tp53^-/-^* mice. Graph representing the mRNA levels of *Trim71* in lung cells isolated from WT and *Tp53^-/-^* mice with PR8 infection. Shown are the individual mice with bar respresenting the mean and error bars represent ± SD. ^∗^ p<0.05 by unpaired t-test. n=1 independent experiments with a total of 5 WT and 8 *Tp53^-/-^* mice. **(H)** same as in **(H)** but for *Ifnb.* ^∗^ p<0.05 by unpaired t-test. **(I)** same as in **(H)** but for *Ifnar2.* ^∗^ p<0.05 by unpaired t-test

To further explore the effects of AGO2 in the nucleus upon viral infection, we infected *Tp53^-/-^* and C57BL/6 WT mice and harvested lungs at 3 d.p.i (**Fig 8D**). Supporting our *in vitro* findings, AGO2 was not able to translocated to the nucleus in *Tp53^-/-^* mice (**Fig 8E**). Moreover, in the *Tp53^-/-^* mice, we observed a significant reduction of viral mRNA levels and infectious virus in the lungs (**Fig 8F,G**). TRIM71 was the second top target from our nuclear fPAR-CLIP and a negative regulator of AGO2 and IFN levels. Interestingly, IAV-infected *Tp53^-/-^* mice showed enhanced *Trim71* levels (**Fig 8H**), in line with all the results obtained *in vitro*. Furthemore, *Tp53^-/-^* mice had increased *Ifnb* and *Ifnar2* mRNA levels (**Fig 8I,J**), again supporting the *in vitro* mechanistic insights. Collectivly, our data indicate that absence of *Tp53 in vivo* significantly reduces viral infection.

To test whether pharmacological intervention could also result in beneficial antiviral effects, we administered the FDA-approved ATO treatment known to monomerize p53 and destabilize nuclear AGO2 (**Fig 3 and Sup Fig 3**). Mice received 0.15mg/kg of ATO for 4 days, starting one day before infection, and lungs were harvested at 3 d.p.i. (**Sup Fig 8H**). Two control groups received either daily vehicle injection or daily ATO injection, without infection. The experimental groups received either daily vehicle injection + IAV i.n. infection or daily ATO injection + IAV i.n. infection. At 3 d.p.i. single cells were isolated from lungs. It is important to note that it was not possible to fully assess the efficacy of the ATO treatment *in vivo*. Indeed, when using nuclear p53 levels, as a proxy of efficacy, ATO treatment was successful in only ∼50% of treated mice. However, whenever p53 was excluded from the nucleus, we observed that also AGO2 was excluded regardless of IAV-infection, underscoring the critical role p53 plays in nuclear AGO2 accumulation also *in vivo* (**Sup Fig 8I**). Therefore, for our subsequent analysis we only considered mice where ATO treatment excluded p53 from the nucleus. Indeed, also in this pharmacological model, we observed a trend towards reduced viral mRNA levels and ∼1-log reduction of infectious virus in the lungs of ATO-treated mice, albeit not significant (**Sup Fig 8J,K**). Furthermore, *Trim71*, *Ifnb* and *Ifnar2* mRNA levels were significantly enhanced in the ATO-treated mice (**Sup Fig 8L-N**).

Taken together, we unraveled a new layer of regulation of IAV infection and propose that targeting either p53 or nuclear AGO2 might serve as a potential therapeutic avenue for IAV modulation.

## DISCUSSION

Unveiling mechanisms of viral resistance is crucial for designing effective new therapies to alleviate disease. Here, by combining classical biochemical fractionation experiments with fPAR-CLIP to identify the precise targets of nuclear AGO2 and *in vivo* mouse models we discovered complex events leading to increased viral replication. We identified viral infection as a potent trigger for nuclear AGO2 translocation, in complex with p53. In the nucleus, AGO2 suppresses innate immune genes thus favoring viral replication. By pinpointing the molecular mechanisms involved, we could use the FDA approved drug arsenic trioxide to reverse AGO2 nuclear localization, increase innate immune response and lower viral infectivity.

In our study, we highlighted how certain viral components—specifically the large T antigen from SV40 (a DNA virus) and NS1 from IAV (an RNA virus)—induce the nuclear accumulation of both p53 and AGO2 (**Fig 2D,E**). This adds to the multiple proviral roles of NS1 ^30^. A limited number of previous reports already identified nuclear presence of AGO2 upon IAV infection ^22,62^, however both utilized cell lines (A549 and HEK293T) which are already nuclear-AGO positive at steady state ^25^, thus complicating the interpretation of the results. Herein, by using HEK293 cells, which are nuclear AGO negative at steady state, we could better mimic what happens in mouse lungs. Indeed, also Wang et al described nuclear accumulation of AGO2, mediated by NS1, and associated with increased virulence *in vivo*, but did not provide any mechanistic explanation of the phenomenon ^62^. In general, the translocation of AGO2 from the cytoplasm to the nucleus is an intricate, dynamic process, elicited by a spectrum of cellular stressors, including, but not limited to, cell confluence, DNA damage, activation of oncogenes, and viral infections ^20,62–65^. In our investigation, we observed nuclear accumulation of AGO2 specifically in response to acute IAV infection (MOI ≥ 2). This phenomenon was concomitant with the elevated expression of *GADD45B* (log2 fold change: 3.129), a key player in DNA damage repair and cellular senescence^66^. Fascinatingly, the nuclear accumulation of p53 is also a characteristic feature of senescent cells ^67^. Many *in vitro* studies on innate antiviral immunity have been performed at low MOI, however in our work we could observe only an effect starting from MOI 2. Interestingly, a detailed study analyzing NS1 expression level and timing elegantly demonstrated that higher MOI is indeed essential for potent and early NS1 expression ^68^. Such expression was negatively correlated with immune-related genes thus suggesting that the number of virions infecting a single cell determined the antiviral response of that specific cell. Thus, we believe that early and potent NS1 induction is a prerequisite for AGO2 nuclear translocation *in vitro* at the time points we have analyzed. Possibly, lower MOI may also induce the same phenomena once NS1 is expressed at higher level, which may take longer, but it was impossible to experimentally assess due to technical limitations (**Sup Fig 1A**). Strongly supporting the physiological relevance of our findings is the remarkable AGO2 nuclear translocation in lungs of IAV infected mice (**Fig 8C**): *in vivo* cells are infected by single virions, initially, but thereafter neighboring cells are infected at high MOI.

In addition, here we demonstrate that p53 is necessary to stabilize AGO2 in the nucleus. We postulate that the tandem tryptophan-binding pockets within the PIWI domain of AGO2 may serve as interaction sites with the flexible N-terminus of p53, characterized by three tryptophan residues: Trp23, Trp53, and Trp91 (**Fig 3C)**. While it was previously reported that, in cancer, p53 and AGO2 interacted ^69^, here we further revealed the strong interaction within the nucleus and complement it by demonstrating that the tetramerization of p53 may enhance the stability of nuclear AGO2 (**Fig 3J**). p53 N-terminus, containing transactivation domains and multiple phosphorylation sites, can also modulate DNA binding, potentially influencing its interaction with proteins like MDM2, a p53 E3 ligase ^70^. Further research is needed to understand how AGO2-p53 interactions impact p53’s DNA binding and protein stability.

Here, we further elucidated the functional consequences of AGO2 nuclear translocation. In general, RNAi function during viral infections is intricate and hotly debated: it has been shown that RNAi can inhibit viral replication and augment the host immune response, thus acting as an anti-viral factor, or promote viral replication and host immune response evasion as a pro-viral factor ^4,7,71,72^. While we have not studied in detail the potential impact of direct RNAi against IAV, we have indirect evidence suggesting this does not play a major role. Indeed, we did not detect neither enrichment of AGO targeting viral genes nor did we detect any vial RNAs being loaded to AGO. Furthermore, by silencing p53 we did not affect overall AGO2 levels, nevertheless we measured differences in viral mRNA, which was not compatible with a direct antiviral role of RNAi. Notably, diminished levels of AGO2 mRNA are reported in COVID-19 patients in comparison to healthy individuals ^73^. Moreover, AGO4, another effector in the RNAi/miRNA pathway, has illustrated antiviral properties in mammals ^74^. Together, the effects of AGO2 on viral infection can be governed by diverse variables such as viral quantity, host cell type, expression level and subcellular localization of AGO2. Thus, a nuanced understanding of the viral infection context is essential to decrypt the varied roles of AGO2 in viral infections.

Our experimental data, combined with analysis of nuclear AGO targets by fPAR-CLIP and miRNA-seq strongly indicated that nuclear AGO2 has a direct role in silencing the antiviral interferon response in infected cells. This finding is consistent and mechanistically explains, several sparse observations from previous studies: Backers *et al*. showed that in the absence of small RNAs, *in vivo* RNA virus infection reached lower titers due to reduced repression of antiviral genes; further, Seo *et al*, also postulated that inactivation of RISC would facilitate antiviral response ^6,7^. Yet another study identified AGO2 as negative regulator of IFNβ signaling and another reported that p53 had direct impact on IFN-regulated genes, without relying on its transcriptional activity ^62,75^. Finally, it was also shown that Dicer-2 accumulation had a negative effect on IFNβ signaling in human cells ^76^. Altogether, the studies summarized provided several pieces of information which we confirmed and expanded here in a comprehensive mechanistical model, which includes a nuclear function of AGO2. Furthermore, IAV NS1 causes global RNA PolII termination defects^77^ and may promote accumulation of aberrant transcripts, targeted by AGO2 in the nucleus. While this may play some role, here we demonstrate that AGO2 is required for the effects observed.

In our quest to understand the molecular details by which IAV promotes AGO2:p53 accumulation within the nucleus, we utilized fPAR-CLIP to scrutinize the silencing targets of nuclear AGO. Beside the IFN-pathways genes, discussed above, we underscored the pivotal roles of nuclear E3 ligases, notably TRIM71 and MDM2. These ligases are instrumental in determining the degradation pathways of AGO2 and p53, thereby influencing their nuclear stability ^54,78^. Our observations also underlined nuclear AGO targeted entities which are crucial for p53 stability and its export to the cytosol, namely MDM2 and XPO1. Together, our results indicate that the virus-induced presence of nuclear AGO2 appears to facilitate the aggregation of AGO2:p53 in the nucleus and thus serving the dual role of stabilizing the complex and to repress the antiviral immune response. TRIM71 was one of the top targets, also validated by multiple miRNA targeting, and, beside its E3 ligase role, it has not been extensively studied; neverthelessm it has been reported as immune enhancing protein, and, in a recent study, its silencing resulted in increased SARS-CoV-2 viral titers ^11,79^. Here we further validated TRIM71 antiviral role and showed it to be associated with viral titer, IFN response and AGO2 stability both *in vitro* and *in vivo*.

The findings from our current work elucidate how the activation of p53, induced by IAV, fosters the nuclear accumulation of AGO2, subsequently leading to the suppression of innate immune genes, a scenario that can aggravate the clinical outcomes of IAV infection. Such insights into the interplay between p53 activation and AGO2 translocation underscore the potential of targeting p53-mediated AGO2 nuclear translocation as a viable therapeutic strategy, as we have demonstrated using arsenic trioxide. Overall, our results could open new avenues to slow down the progression and reduce the severity of viral infections.

## LIMITATIONS OF THE STUDY

Here we demonstrated a link between nuclear AGO2:p53 translocation, the suppression of IFN response and viral loads upon IAV infection. Although likely, we can not generalize our results to other viral pathogens and/or other disease models, like cancers, which show increase accumulation of nuclear AGO2. In addition, while we have individually silenced p53 and AGO2 and shown that the majority of the effects we observe are due to AGO2, we cannot fully exclude that stabilized nuclear p53 will impact transcription thus influencing some of our results. Furthermore, broad changes in transcription upon *TP53* deletion may also contribute to viral resistance in cell lines and mice and should be investigated in future studies.

## ACKNOWLEDGMENTS

This work was funded by the European Research Council (ERC-StG, B-DOMINANCE, grant no. 850638 to DA); the Swedish Research Council (grant no. 2021-01164, 2021-01165 to DA and grant number 2019-01855 to AAS); the Knut and Alice Wallenberg Foundation (grant no 2021.0033 to DA and grant no PAR 2020/228 to AAS); the Swedish Society for Medical Research (grant no S19-0019 to AAS) and the University of Gothenburg. J.O.W. is financially supported by the Knut and Alice Wallenberg Foundation as part of the National Bioinformatics Infrastructure Sweden at SciLifeLab. We are grateful to Dr. Marianne Farnebo (Karolinska Institutet) for the generous contribution of TP53L expressing MCF7 cells. We would like to thank the staff at the Experimental Biomedicine (EBM) core facility at the University of Gothenburg for animal management. We thank SciLifeLab CRISPR Functional Genomics unit at Karolinska Institutet for generating the TP53 KO HEK293 cells. The sequencing was performed at the National Genomics Infrastructure (NGI).

## AUTHOR CONTRIBUTIONS

H.-C.H. designed the experiments, performed the experiments, analyzed data, and wrote the manuscript. D.A., D.G.A, and J.O.W. analyzed the NGS sequencing data. I.N. generated the PAR-CLIP libraries, miRNAseq libraries and Let-7 qPCRs data. V.L. generated the PAR-CLIP libraries and miRNAseq libraries. D.F.B. generated mutant NS1 virus. C.F. performed immunofluorescent experiments. K.S., A.A.H.P., and C.W. performed animal experiments and analyzed data. V.I.S. provided *Tp53*^-/-^ mice, scientific input and feedback on the manuscript. D.A. and A.A.S. supervised the entire project, designed the experiments, analyzed data, and wrote the manuscript. All authors contributed to the manuscript

## METHODS

### Mice and Ethical Statement

All the experiments were conducted according to the protocols (Ethical permit numbers 1666/19, 38/23 and 2071/19) approved by regional animal ethics committee in Gothenburg. Female, 8-12 weeks old C57BL/6 mice and *Tp53*^-/-^ mice were purchased from Janvier, France. They were housed in the specific pathogen-free animal facility of Experimental Biomedicine Unit at the University of Gothenburg.

### Cell culture

HEK293 (ATCC, CRL-1573), A549 (ATCC, CCL-185), HEK293T (ATCC, ACS-4500), MCF7 (ATCC, HTB-22), MCF7 TP53L (kind gift from Dr. Marianne Farnebo, Karolinska Institutet) and MDCK (ATCC, CCL-34) cell lines were cultured in Dulbecco’s modified Eagle’s medium (DMEM) (Gibco, 11995065), supplemented with 10% fetal bovine serum (FBS) (Gibco, 11573397) and 100 U/ml penicillin-streptomycin (Gibco, 11548876) in a humidified incubator at 37℃ and 5% CO2. The SK-N-BE(2) cell line was cultured in the same medium, but with the addition of 1% non-essential amino acids (Gibco, 11140050). Vero cells (kind gift from Dr. Kristina Nyström, University of Gothnburg) and MDCK cells, were grown in DMEM (Gibco, 11594446) supplemented with 10% FBS (Gibco, 11550356) and 10 μg/ml gentamicin (Gibco, 15710064) at 37℃. For the propagation of rescued virus in Vero cells, DMEM was supplemented with 0.3% BSA (Sigma, A7906), 1 μg/ml TPCK-trypsin (BioNordika, LS003740) and 10 μg/ml gentamicin. For the infection of Vero cells with the rescued virus, DMEM supplemented with 1 mM HEPES (Gibco, 11560496), 5 μg/ml gentamycin and 1 μg/ml TPCK-trypsin was used.

### TP53 knockout HEK293 by CRISPR

Single guide (sg)RNAs were formed by duplexing crRNAs (5’ GCAGTCACAGCACATGACGG-3’ and 5’-AATCAACCCACAGCTGCACA-3’; sg1 and sg2 respectively; Alt-R® CRISPR-Cas9 crRNAs; IDT) with Alt-R™ CRISPR-Cas9 tracrRNA, ATTO™ 647 (IDT) according to manufacturer’s instructions. Equimolar mixtures of the two sgRNAs were precomplexed with Cas9 protein (Alt-R® S.p. Cas9 Nuclease V3, IDT) into ribonucleoproteins (RNPs). RNPs were introduced by electroporation into HEK293 cells using the Neon system (ThermoFisher Scientific). Cells were expanded post-electroporation for several days prior to genomic DNA extraction (QIAamp mini kit, QIAGEN).The frequency of edited alleles was in first instance estimated by droplet digital PCR (ddPCR, QX200 System, BioRad) in a dropoff assay, using a reference probe combined with a probe specific for the wild type allele. After single-cell isolation, successful p53 knockout was confirmed by western blot analysis, showing complete loss of p53 protein.

### Whole cell lysate and biochemical fractionation

To extract whole cell lysates, the cells were first washed with cold 1xPBS and then lysed on ice with RIPA lysis buffer (50 mM Tris-HCl, pH 7.6, 150 mM NaCl, 1 mM EDTA, 1% NP-40, 1% sodium deoxycholate, 0.1% sodium dodecyl sulfate) supplemented with a protease inhibitor cocktail (Merck Millipore, 04693132001). Samples were cleared with centrifugation at 12 000 *g* for 20 minutes and the supernatant collected. Biochemical fractionation assay was done as previously described ^25,80^. Briefly, cell pellets were gently dissolved in a hypotonic lysis buffer (10 mM Tris-HCl, pH 7.6, 10 mM NaCl, 3 mM MgCl_2_, 0.3% NP-40, 10% Glycerol), supplemented with a protease inhibitor cocktail, with gentle pipetting up and down and collected with centrifugation for 2 minutes at 200 g. The supernatant was cleared with centrifugation at 12 000 *g* for 20 minutes and the supernatant stored as the cytoplasmic fraction. The remaining nuclear pellet was washed 3 times with the hypotonic lysis buffer and each time collected by centrifugation for 2 minutes at 200 g. Each time the supernatant was discarded. From the remaining pellet, the nuclear proteins were extracted using a nuclear lysis buffer (20 mM Tris-HCl, pH 7.6, 150 mM KCl, 3 mM MgCl_2_, 0.3% NP-40, 10% Glycerol) supplemented with protease inhibitor cocktail. The lysate was sonicated twice for 10 seconds each time at 60% amplitude (Sonics, VCX130). The nuclear fraction was cleared with centrifugation at 12 000 *g* for 20 minutes and the supernatant collected. Protein concentration was measured using Bradford Reagent (B6916, Sigma Aldrich).

### Western blotting

After protein extraction, as described above, 5-20 ug of protein were used for western blot experiments. Protein samples were run on 4-12% Bis-Tris gels and transferred onto Nitrocellulose membrane (Cytiva, 1060000). Proteins of interested were analyzed by hybridization with their corresponding antibodies (see below) and visualized by chemiluminescence using Thermo Scientific SuperSignal™ West Dura Extended Duration Substrate (ThermoFisher, 34076).

|  |
| --- |
| FLAG (Merck, M8823) |
| HA (Sigma, H9658) |
| AGO1 (CST, 5053S) |
| AGO2 (Abcam, ab32381) |
| AGO2 11A9 for IF (Merck, MABE253) |
| AGO3 (CST, 5054) |
| β-Tubulin (Proteintech, 66240-1-Ig) |
| YB1 (Abcam, ab12148) |
| Histone H3 (CST, 44995) |
| p53 (Abcam, ab1101) |
| p53 (Santa cruz, sc-126) |
| p53 (ThermoFisher, MA5-11296) |
| GAPDH (Abcam, ab9485) |
| SV40 (Santa cruz, sc-147) |
| NS1 (ThermoFisher, PA5-32243) |
| TNRC6A (GW182) (Santa Cruz, sc56314) |
| mCherry (Santa cruz, sc-101529) |
| Ubiquitin (Santa cruz, sc-8017). |
| TRIM71 (ThermoFisher, PA5-19281) |
| IFNAR2 (ThermoFisher, PA5-28303) |
| LIN28A (Santa cruz, sc-293120) |
| LIN28B (ThermoFisher, PA5-50609) |
| anti-rabbit IgG, HRP conjugate (Sigma, GENA934) |
| anti-mouse IgG, HRP conjugate (Sigma, GENA931) |
| anti-goat IgG, HRP conjugate (ThermoFisher, 10466033) |
| anti-sheep IgG, HRP conjugate (Merck, AP147P) |
| Anti-mouse Fc BP-HRP (Santa Cruz, sc525416) |
| Anti-rabbit IgG (Conformation Specific) (L27A9) mAb (HRP Conjugate) #5127 |

### Plasmid transfection

Plasmids encoding the Influenza A virus components PB1, PB2, NP, M, HA, NA, NS1, PA and mCherry-NS1 were a kind gift from Dr. Ivan Kosik (NIH, USA). The SV40 Large T antigen and HA-TRIM71 were purchased from Addgene (plasmid # 136616 and #52717, respectively). LIN28A and LIN28B were also purchased from Addgene (plasmid # 51387 and #51373, respectively). 3 µg of plasmid was used for transient transfection using X-tremeGENE™ HP DNA Transfection Reagentreagent (Roche, 6366236001) following the manufacturer’s instructions.

### NS1 mutagenesis

The pDZ plasmid encoding NS1, kindly provided by Dr. Ivan Kosik from the NIH, USA, was used as a template for site-directed deletion of its C-terminal regions spanning amino acids 81-225 and 125-225. The resulting constructs encoded truncated NS1 proteins comprising amino acids 1-80 and 1-124, respectively. These constructs were transiently transfected into HEK293 cells using the X-tremeGENE HP DNA Transfection Reagent (Roche, 6366236001) according to the manufacturer’s instructions. The expression of both full-length and truncated NS1 mutants was confirmed by western blotting using an anti-NS1 antibody (ThermoFisher, PA5-32243).

### Viral infection in cells

1 million HEK293 cells were seeded in 10 cm dishes. Cells were allowed to attach for 8 hrs and were infected with PR8 with different MOI (0.1 to 10) for 16 hours in serum free media. Cells were collected by trypsinization.

### Viral titer determination from infected lungs

MDCK cells were seeded at 50,000 cells/well in 96-well plates. After overnight incubation, cells were washed twice with PBS. Harvested lungs were placed in PBS at a constant w/v ratio. Homogenized lungs were 10-fold diluted starting 1:10 in infection media (DMEM containing 0.1% BSA (fraction V; Roche), 10 mM HEPES (Corning), 500 µg/ml gentamicin (Gibco), and 1 µg/ml TPCK trypsin (Worthington) and incubated on MDCK cells. After 3 days cytopathic effect was visualized after crystal violet staining and TCID_50_ titer was calculated using the Spearman and Karber method.

### Immunofluorescence staining and microscopy

To perform immunofluorescence assay, HEK293 cells were either transfected with mCherry-NS1 or infected with PR8-IAV, as described above, for 16 hours. Alternatively, HEK293 cells were infected with mCherry-PR8 for 16 hours. Cells were washed with PBS and fixed with 4% buffered formalin for 15 minutes at room temperature. Cells were then washed twice with PBS and blocked with 5% BSA in PBS for 1 hour at room temperature. Finally, cells were incubated with 1:200 AGO2 (Merck, MABE253) overnight. The next day cells were washed 3x with PBS and probed with secondary antibodies: 1:2000 Alexa Fluor® 488 Goat Anti-Rat IgG (Invitrogen, 10729174) or 1:1000 phalloidin (Invitrogen, 10643853) for 2 hours at room temperature. To visualize the cell nuclei, 4’,6-diamidino-2-phenylindole (DAPI; Invitrogen, D3571) was added for 5 minutes in the dark. Slides were mounted using 10 μl Prolong Diamont (Invitrogen, 15372192). Confocal images were taken on a Zeiss LSM780 and the images were analyzed using ImageJ® software and Affinity Designer®.

### Doxorubicin treatment

Doxorubicin (Biotechne (Tocris), 2252) was dissolved in dimethyl sulfoxide (DMSO) to prepare a 1 mM stock solution. The stock solutions were stored at −20℃ and diluted in the culture medium to 1 µM final concentrations. Cells were incubated with doxorubicin for 24 hours.

### Immunoprecipitation assays and AGO protein affinity purification with T6B peptide

Immunoprecipitation was carried out in 1-3 mg of protein lysate and 2 μg of anti-AGO2, anti-IgG, or anti-p53 antibodies. For purification of AGO1-4, 400 μg Flag-tagged T6B peptide was used ^50^. Dynabead Protein G beads (10004D, ThermoFisher) or anti-Flag M2 beads (M8823, Millipore) were conjugated with either antibodies or T6B peptide, respectively, for 4 h, washed and incubated with protein lysates. Next, beads were washed three times with RIPA buffer and bound proteins were eluted at 95°C for 5 min in 3× SDS Laemmli buffer and assessed by western blot. The pull-down efficiency was confirmed by western blot.

### p53 mutagenesis

The plasmid encoding amino acids 1-393 of the p53 protein with a FLAG tag at the C-terminus (Addgene plasmid #10838) was used as a template for site-directed mutagenesis to delete the N-terminal regions spanning amino acids 1-31, 1-62, or 1-93. The resulting PCR products were cloned into the pcDNA3.1 expression vector (Invitrogen, v79020). The final constructs encoded truncated p53 proteins with a FLAG tag starting at amino acids 32, 63, or 94, respectively. The pcDNA3 constructs were used for transient transfection in HEK293 cells using an X-tremeGENE HP DNA Transfection Reagent (Roche, 6366236001) according to the manufacturer’s instructions. The expression of the full-length and truncated p53 proteins were verified by western blotting using an anti-FLAG antibody (Merck, M8823).

### Arsenic trioxide (ATO) treatment

Arsenic trioxide (Merck, 202673-5G) was dissolved to 100 mg/ml stock solution in NAOH which was further diluted in DMEM to 1 mg/ml. The stock solutions were stored at −20℃ and diluted in the culture medium to 0.01, 0.1 or 0.5 μg/ml concentrations before use. Cells were incubated with ATO for 24 hours.

### siRNA Gene Silencing of AGO2 and TP53

Small interfering RNA (siRNA) targeting human *AGO2* (siRNA ID: s109013 and ID s25931), *TP53* (siRNA ID s607), and scramble control siRNAs (siRNA ID: 4390843) were purchased from ThermoFisher Scientific. Human A549, HEK293T, MCF7, or SK-N-BE(2) cells were transfected with the siRNAs using Lipofectamine RNAiMAX Transfection Reagent (ThermoFisher Scientific, #13778030) according to the manufacturer’s instructions. The cells were harvested 48 hours after siRNA transfection to evaluate AGO2 or p53 knock-down efficiency by quantitative RT-qPCR and protein level by western blotting.

### MG132 treatment and ubiquitination assay

MG132 (ThermoFisher, 15465519) were dissolved in dimethyl sulfoxide (DMSO) to prepare a 10 mM stock solution. The stock solutions were stored at −20℃ and diluted in the culture medium to 40 μM concentrations before use. Cells were treated with MG132 or DMSO only for 4 hours. Following the treatment, cells were lysed and/or fractionated and assessed by western blotting.

### Protein-protein docking and structure modelling

To predict the interaction between PIWI domain of AGO2 (570-859 amino acid) with the T6B peptide from TCRC6B (599-683 amino acid) or N-terminal of p53 (1-94), we performed molecular docking by using the HDOCK server ^42^. This server uses the hybrid algorithm of template bases modeling and ab initio free docking and provides the top ten complex models with the highest scores. Among the top 10 models for both complexes, the first models with the lowest docking scores and highest confidence scores. Specifically, the first model of the AGO2 with T6B peptides from TNRC6B complex has a docking score of −308.41 and a confidence score of 0.9596, while the first model of AGO2 with N-terminal of p53 complex has a docking score of −268.89 and a confidence score of 0.9151. To model the structures of p53 (AF-P04637-F1), we downloaded their atomic coordinates from the AlphaFold2 database ^81^. We used the PyMOL Molecular Graphics System, Version 2.3.4 (Schrödinger, LLC, https://pymol.org/2/) to visualize and modify the structure figures.

### Real-time quantitative PCR (RT-qPCR)

To analyze gene expression levels, real-time quantitative PCR (RT-qPCR) was performed on the following cell lines: HEK293, TP53 KO HEK293, HEK293T, A549, MCF7 and MCF7 TP53L. Total RNA was isolated from each cell line using the Quick-RNA Miniprep Kit (ZYMO Research, R1055) following the manufacturer’s instructions. To generate cDNA, 1 μg of total RNA was used in a reverse transcription reaction with the iScript cDNA Synthesis kit (Bio-Rad, 1708891) according to the manufacturer’s instructions. The RT-qPCR reactions were performed in a 10 μL mixture, consisting of 1x iQ™ SYBR® Green supermix (Bio-Rad, 1708880), 0.5 μmol/L of each primer, and 10 ng of cDNA template. The RT-qPCR result was acquired by CFX Connect Real-Time PCR Detection System (Bio-Rad) using the following primers:

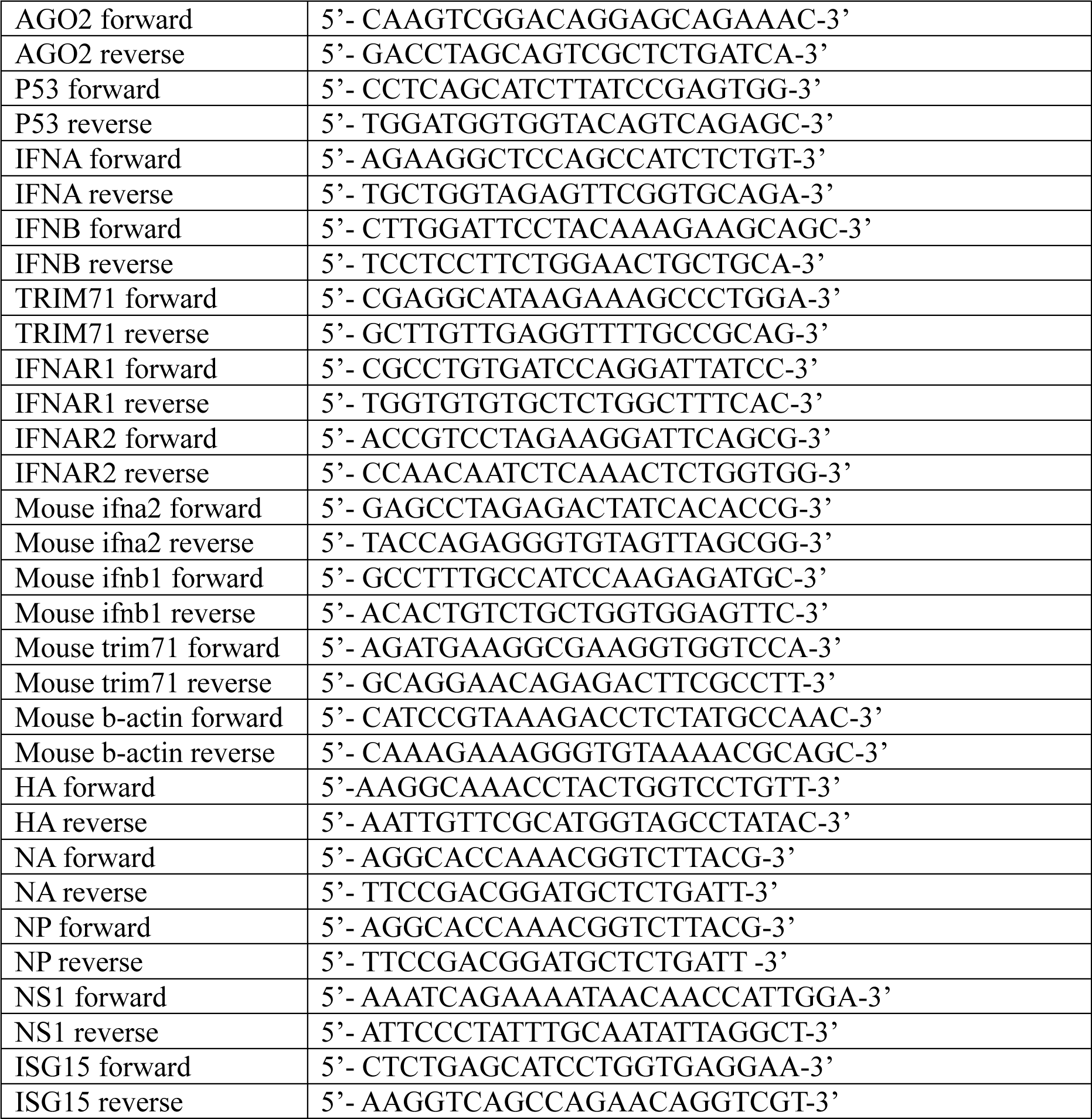
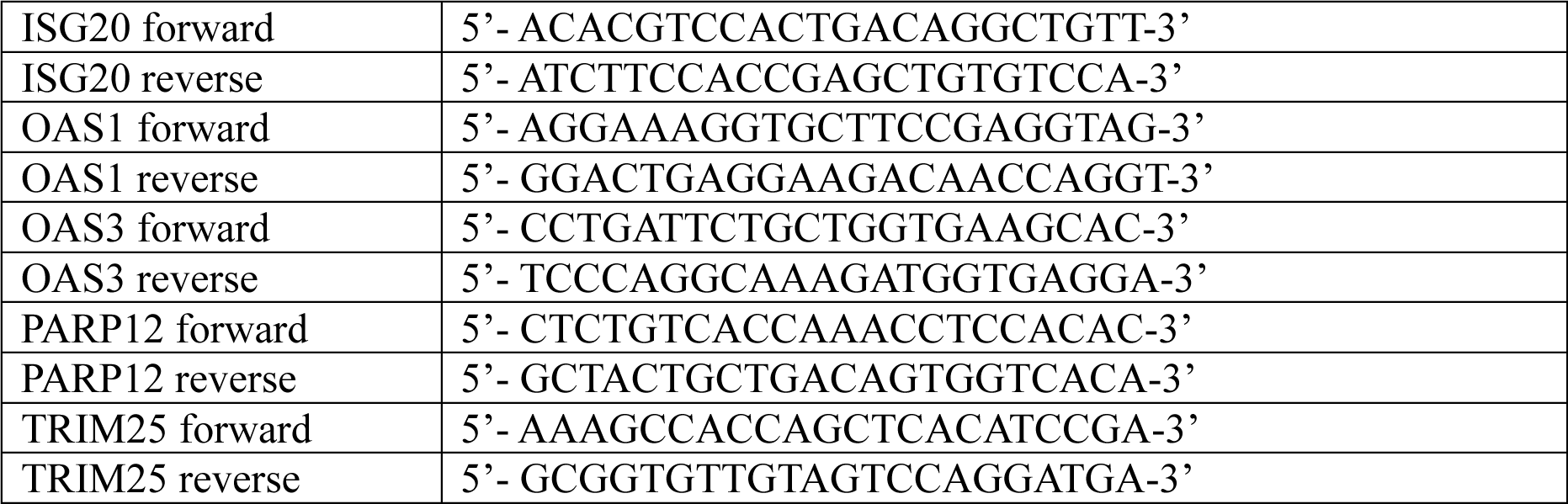

### Flow cytometry

HEK293 and TP53 KO HEK293, as well as MCF7 and MCF7 TP53L cells were infected with PR8-NS1-mCherry at multiplicity of infection (MOI) 10 for 16 hours. Cells were collected by trypsinization and resuspended in FACS buffer. Cells were acquired using BD LSR II flow cytometer (BD Bioscience) to measure the expression of mCherry in HEK293 and TP53 KO HEK293, as well as MCF7 and MCF7 TP53L cells. Data analysis was performed using FlowJo V10 software (Treestar).

### Luciferase assay

Cells were seeded 200,000 cells in 12 well dishes. The next day, the cells were transfected with ISRE reporter plasmid (a gift from Dr. Chia-Wei. Li, Academia Sinica, Taiwan) or with the internal control SV40 renilla plasmid (Promega) for 24 hours and subsequently infected with PR8 at MOI 10 for an additional 16 hours. Cells were then collected for luciferase assay using the Dual-Luciferase Reporter assay (Promega, E1910), following manufacturerś instructions. Plates were read using CLARIOstar Plate Reader (BMG Labtech).

### RNA-sequencing

Total RNA was extracted from HEK293 with or without PR8 infection (MOI:10) for 16 hours, using the Quick-RNA Miniprep Kit (ZYMO Research) following the manufacturer’s protocol. the concentration and quality of the RNA was analyzed using Agilent 2200 TapeStation System. RNA samples with RNA Integrity Number higher than 8 were sent to SNP&SEQ Technology Platform (NGI Uppsala, Sweden). Libraries were prepared from 300 ng RNA using the Illumina Stranded Total RNA library preparation kit, including Ribo-Zero Plus treatment (20040525/20040529, Illumina Inc.) according to manufacturer’s instructions. For indexing Unique Dual Indexes (20040553/20040554, Illumina Inc.) were used. Sequencing was carried out with NovaSeq 6000 system using paired-end 150 bp read length, S4 flowcell and v1.5 sequencing chemistry. As a control sequencing library for the phage PhiX was included and a 1% spike-in in the sequencing run. RNAseq data were preprocessed using the RNAseq nf-core pipeline ^82^. Differential expression analysis was done using DEseq2 ^83^, on genes with at least 10 reads in at least 3 samples. Genes with FDR adjusted p-value < 0.01 and absolute log2 fold change > 0.5 were considered differentially expressed. Hypergeometric tests, implemented in TopGO, were used to look for enriched Gene Ontology annotation among the differentially expressed genes. The fraction of reads mapping to introns and other genomic regions was calculated using ReSQC ^84^.

### Fluorescent PhotoActivatable Ribonucleoside-enhanced CrossLinking and ImmunoPrecipitation (fPAR-CLIP)

AGO fPAR-CLIP was carried out by isolating the proteins using the T6B peptide as mentioned above. fPAR-CLIP library preparation, sequencing and initial data processing was performed as described in ^25,49^ with minor modifications. Briefly, to obtain AGO proteins RNA footprints, unprotected RNA was digested on beads with 1 U RNase T1 (EN0541, ThermoFisher) for 15 min at RT. Next, the beads were washed three times with RIPA buffer and three times with dephosphorylation buffer (50 mM Tris–HCl, pH 7.5, 100 mM NaCl, 10 mM MgCl_2_). After washing, the protein-bound RNA was dephosphorylated with Quick CIP (M0525S, New England Biolabs) for 10 min in 37°C. Post dephosphorylation the beads were washed three times with dephosphorylation buffer and three times with PNK/ligation buffer (50 mM Tris-HCl, pH 7.5, 10 mM MgCl_2_). Following, 0,5 μM fluorescently tagged 3’ adapter (MultiplexDX) were ligated with T4 Rnl2(1–249)K227Q (M0351, New England Biolabs) overnight at 4°C and washed three times with PNK/ligation buffer. Next, RNA footprints were phosphorylated using T4 PNK (NEB, M0201S) for 30 min in 37°C and washed three times in RIPA buffer. To release the proteins the beads were incubated at 95°C for 5 min in 3× SDS Laemmli buffer. Next, the eluates were separated on a 4-12% SDS/PAGE gels (NW04122BOX, Invitrogen) and AGO:RNA complexes visualized on the IR 680 channel (Chemidoc MP system, Bio-Rad). Subsequently, appropriate AGO:RNA bands were excised from the gel and protein digested with Proteinase K (RPROTK-RO, Sigma Aldrich) and released RNA isolated via phenol:chloroform phase separation. Following, 5’ adapter ligation (MultiplexDX) was performed on the purified RNA samples with 0,5 μM of the adapter and Rnl1 T4 RNA ligase (ThermoFisher, EL0021) for 1 h at 37°C. Next, the RNA was reverse transcribed using SuperScript IV Reverse Transcriptase (ThermoFisher, 18090010) according to manufacturer’s instructions. The libraries were amplified in a series of PCR reactions performed using Platinum Taq DNA polymerase (ThermoFisher, 10966034) and size selected with 3% Pippin Prep (Sage Science, CSD3010). Sequencing of the libraries was carried out on Illumina NovaSeq 6000 platform. For data processing Bcl2fastq (v2.20.0), Cutadapt (cutadapt 1.15 with Python 3.6.4) ^85^, PARpipe (https://github.com/ohlerlab/PARpipe) and Paralyzer ^86^ were used. The 3’ and 5’ adaptor sequences and sequencing primers used in the study are listed below. For each target gene, the normalized PAR-CLIP signal was calculated as nr reads with T->C conversions / (total number of PAR-CLIP reads * average TPM for target gene * 1e-6)

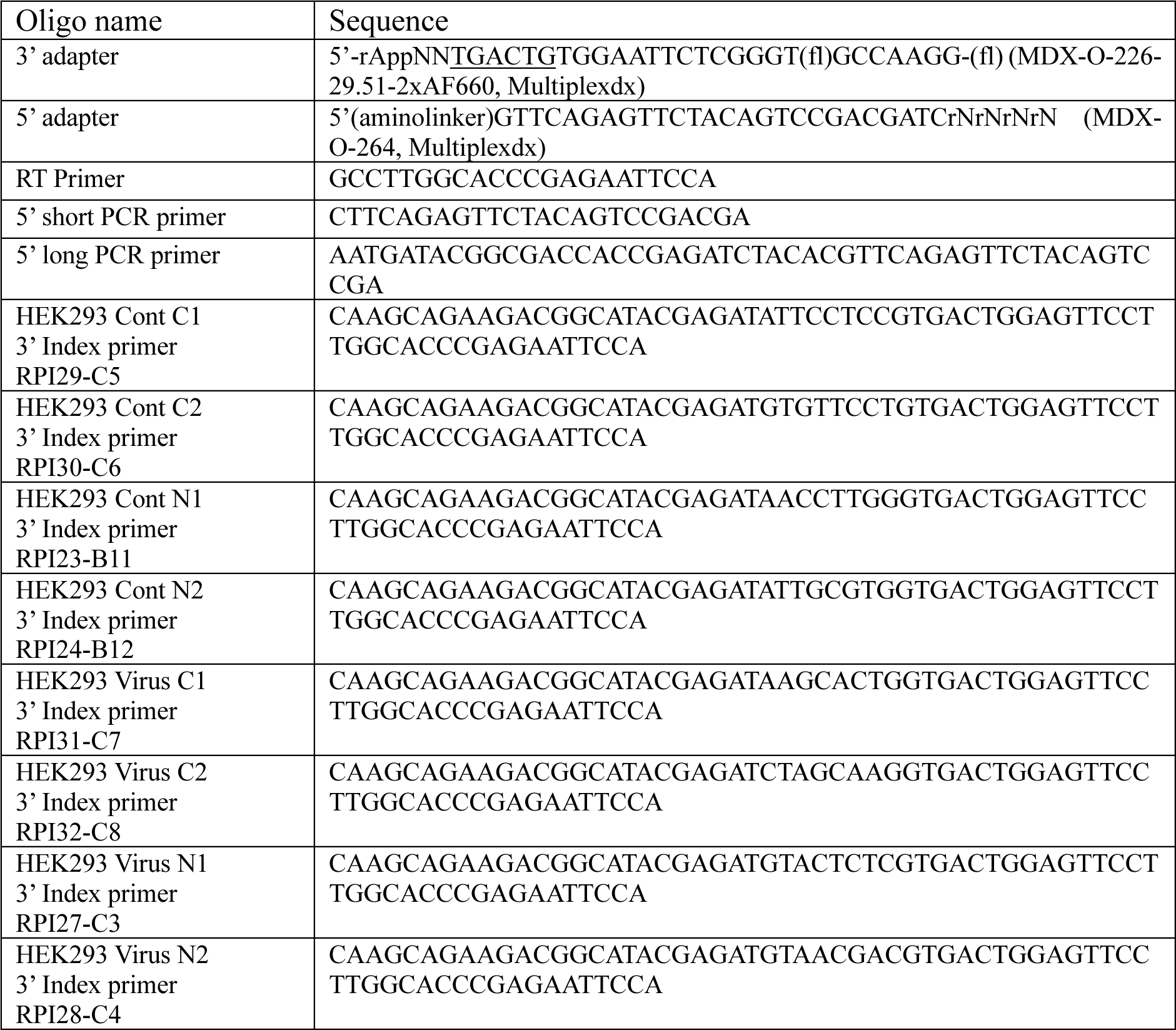

### miRNA sequencing

AGO proteins were immunoprecipitated from 200 mg of protein using Flag-tagged T6B peptide. AGO-bound RNA was recovered from the beads using TRIzol reagent (Invitrogen, 15596026) according to the manufactures instructions and small RNA libraries were produced as previously described ^49^, with minor modifications. Briefly, 3’ adapters with 5’-adenylated RNA adapter (see 3’ adapters in table below) were ligated to the recovered small RNAs using Rnl2(1-249)K227Q RNA ligase (New England Biolabs, M0351) at 4°C overnight with constant shaking. Ligated RNA was pooled within conditions and purified using oligo clean and concentrate kit (ZYMO Research, D4060). Next, the RNA was subjected to 5’ adapter ligation with a 5’ chimeric DNA-RNA adapter (5’aminolinker-GTTCAGAGTTCTACAGTCCGACGATCrNrNrNrN) using RNA ligase (ThermoFisher Scientific, EL0021) at 37°C for 1 hour. Next, the RNA was purified using oligo clean and concentrate kit and reverse transcribed using SuperScript® IV (ThermoFisher Scientific, 18090010) using RT primer (GCCTTGGCACCCGAGAATTCCA). The cDNA was amplified using Platinum Taq DNA Polymerase (ThermoFisher Scientific, 10966034), according to the manufacturer’s instructions using 5’-medium PCR primer (CTCTACACGTTCAGAGTTCTACAGTCC) and 3’ medium PCR primer (CCTGGAGTTCCTTGGCACCCGAGAATT) for 6 cycles. Then the PCR product was purified using the oligo clean and concentrate kit, eluted with 32 µl of nuclease free water, and size selected (74-88 bp) using 3% agarose Pippin Prep (Sage Science, CSD3010). Following size selection, a second round of (X cycle) PCR was performed using the same polymerase, a 5’-long PCR primer: AATGATACGGCGACCACCGAGATCTACACGTTCAGAGTTCTACAGTCCGA, and 3’ indexed primer (see 3’ index primers in table below). Libraries were sequenced on an Illumina NovaSeq6000. Bcl files were converted to fastq files using bcl2fastq. Adapters were trimmed using cutadapt v 2.4. and reads were mapped to the human miRNAs using bowtie ^87^.

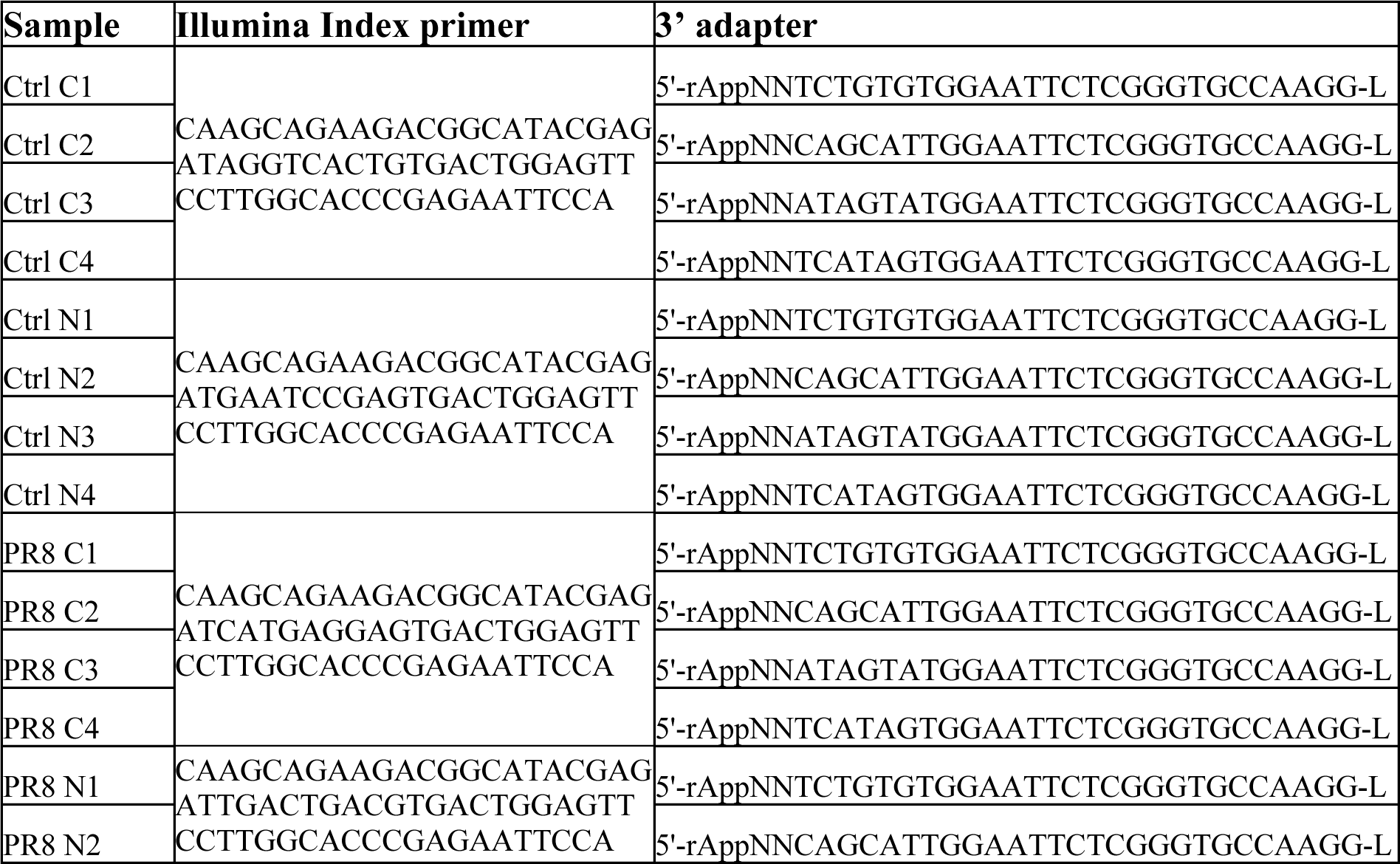
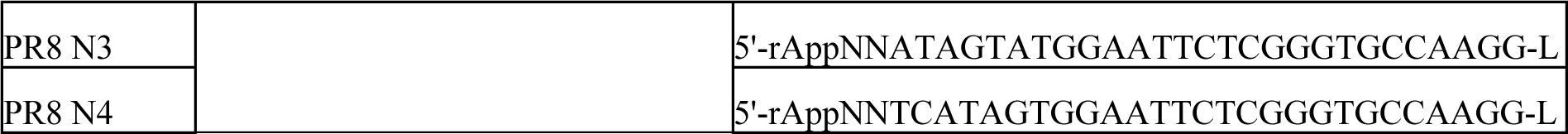

### miRNA RT-qPCR

Reverse transcription of miRNA species was performed with miRCURY LNA™ Universal RT microRNA PCR (Qiagen, 339340). 500 ng total RNA was used per reaction. For miRNA RT-qPCR experiments miRCURY LNA SYBR Green PCR Kit (Qiagen, 339345,) was used. For qPCR reactions, the cDNA was diluted 60 times and miRNA-16-5p was used as the reference gene. Primers for miRNA-16-5p (YP00205702), let-7c-5p (YP00204767), let-7f-5p (YP00204359) and let-7g-5p (YP00204565) were provided by Qiagen. All RT-qPCR experiments were performed in triplicate at least three times, and relative let-7 family miRNAs expression was calculated using the ΔΔCq method ^88^.

### Virus Rescue

Virus containing mutant NS1 genes were rescued as previously described ^89^. Briefly, the day before transfection, HEK293T cells were seeded at 500,000 per well in 6-well plates. The next day, cells were transfected with 1 μg of each plasmid of the eight gene segments of IAV using Lipofectamine 2000 transfection reagent (Invitrogen, 11668-019) in Opti-MEM reduced serum medium (Gibco, 31985-062), according to manufacturer’s instructions. The plates were incubated for 24-, 36-, and 48-hours, and supernatants collected to be used for inoculation of 7 days embryonated chicken eggs and Vero cells for propagation. The presence of the mutant NS1 gene segment after transfection was verified using PCR amplification: Forward primer full length NS1: TGGATCCAAACACTGTGTCAAGC, Reverse primer full length NS1: ACCTAATTGTTCCCGCCATTTCTC), Reverse primer mutant NS1 1-80: TTTCAGAATCCGCTCCACTATCTGC, Reverse primer mutant NS1 1-124: GTCCATTCTGATACAAAGAGGGCCT. Rescue was confirmed by hemagglutination assay.

### Rescued virus TCID_50_ determination

After rescuing the virus, viral titre was assessed using ELISA assay. The day before the assay, 96-well flat-bottomed plates were seeded with 100 μl of 100,000 Vero cells per well and incubated at 37°C overnight to allow for the cells to reach full confluency. Next day, the cells were washed twice with PBS and 180 ml of infection media was added per well. The virus was then added to column 1, at 1:10 dilution in quadruplicate, and ten-fold serially diluted across the plate with the last column as the cells only control. The plates were incubated at 37°C for 20 hours. After incubation, the cells were washed once with PBS and fixed with 50 μl/well ice-cold methanol at 4°C for 10 minutes. Following the fixation, the cells were again washed with PBS then 100 μl/well of the primary ascites anti-NP HB65 antibody (kind gift from Dr. Jonathan Yewdell, NIH), at 1:10000 dilution, was added and allowed to incubate for 2 hours at room temperature. Plates were washed thrice with PBS + 0.05% Tween followed by the addition of 50 μl/well of the secondary rat anti-mouse kappa HRP antibody (Southern Biotech, 1170-05, at 1:1000 dilution, and incubated for 1 hour at room temperature. After a final three times wash with PBS-T, the plates were developed by adding 50 μl/well of TMB (ThermoFisher, 34029) and incubated in the dark for 5 minutes at room temperature. The reaction was stopped with the addition of 25 μl/well of 2M H_2_SO_4_ and the absorbance were read with TECAN Sunrise absorbance microplate reader (16039400) at 450nm. Analysis of the results was carried out using the Reed and Muench infectivity calculator.

### Virus infection in mice

The H1N1 strain of influenza A/PR8 (Puerto Rico/8/34) was propagated in the allantoic cavities of SPF embryonated chicken eggs for 48-72 hours at 37°C. The resulting allantoic fluids were collected, aliquoted, and stored at −80°C until use. Virus titers were assessed by TCID_50_ assay on MDCK cells as previously reported ^90^. To infect the mice, they were anaesthetized with isoflurane, and intranasally inoculated with 2000 TCID50 PR8 in 25 μl sterile PBS/0.1%BSA. Control mice received the same volume of PBS intranasally as a mock infection. After three days of infection, lung tissue samples were collected to isolate single cells using the Lung Dissociation Kit (130-095-927, Miltenyl Biotec, Bergisch Gladbach, Germany) following the manufacturer’s protocol. RNA and protein were extracted from the isolated single cells to determine viral mRNA and AGO2 distribution. Specifically, the RNA was extracted using a Quick-RNA Miniprep Kit (ZYMO Research) following the manufacturer’s protocol, and the protein was extracted using biochemical fractionation method described above. For ATO treatment, mice were injected intraperitoneally with a daily dose of 0.15mg/kg ATO in PBS in a volume of 100 μl for 4 days.

### Data availability

The raw RNA-Seq and PAR-CLIP data described in this paper are accessible through the GEO database (https://www.ncbi.nlm.nih.gov/geo/) under accession no. xxx and xxx. miRNAseq data are accessible through the GEO database (https://www.ncbi.nlm.nih.gov/geo/) under accession no. xxx

### Statistical analysis

The data from three individual experiments waere assessed by unpaired t-test or Mann-Whithney U-test (GraphPad Prism Software Inc, San Diego, CA, USA) and presented as mean ± SD (standard deviation). A p-value < 0.05 was considered statistically significant. Cumulative distribution was analyzed using Kolmogorov-Smirnov Test using the R package stats.

