## Supplemental material for "Nuclear RNAi Modulates Influenza A Virus Infectivity By Downregulating Type-I Interferon Response"

**Supplementary Tables**

**Supplemental Table 1, related to main figure 2.** List of AGO2 associated proteins retrieved from the Harmonizome database

**Supplemental Table 2, related to main figure 3.** Docking models of the PIWI domain of AGO2 with the T6B regions of TNRC6B or the N-terminal region of p53.

**Supplemental Table 3, related to main figure 6.** List of differentially expressed genes from the RNAseq analysis between PR8 infected and non-infected HEK293 cells.

**Supplemental Table 4, related to main figure 6.** Ranked fPAR-CLIP targets in control and PR8-infected, nuclear and cytoplasmic fractions in HEK293 cells.

**Supplemental Table 5, related to main figure 7.** miRNA sequencing from AGO1-4 immunoprecipitation using the T6B peptide from cytoplasmic and nuclear fractions of HEK293 cells with or without PR8 viral infection at MOI 10 for 16 hours.

### Supplementary Figures

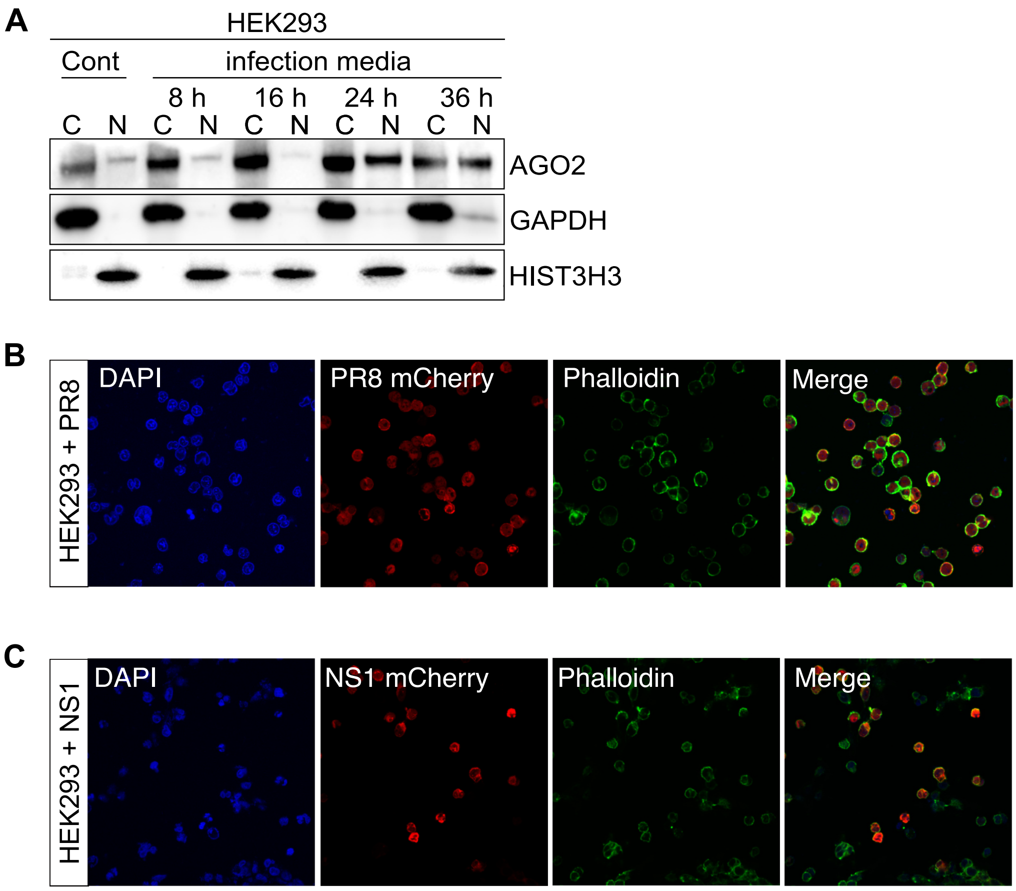


**Supplemental figure 1, related to main figure 1**

**(A)** Representative AGO2 immunoblots from cytoplasmic (C) and nuclear (N) lysates in HEK293 cells. Cells were grown in infection media for 8-36 hours. GAPDH served as cytoplasmic marker and HIST3H3 as nuclear marker. n=3

**(B)** Immunofluorescence images of AGO2 and PR8-mCherry in HEK293 infected with PR8-NS1-mCherry virus at MOI 10 for 16 hours. Phalloidin stained for F-actin and DAPI stained for DNA.

**(C)** Immunofluorescence images of AGO2 and NS1-mCherry in HEK293 infected with PR8-NS1-mCherry virus at MOI 10 for 16 hours. Phalloidin stained for F-actin and DAPI stained for DNA.

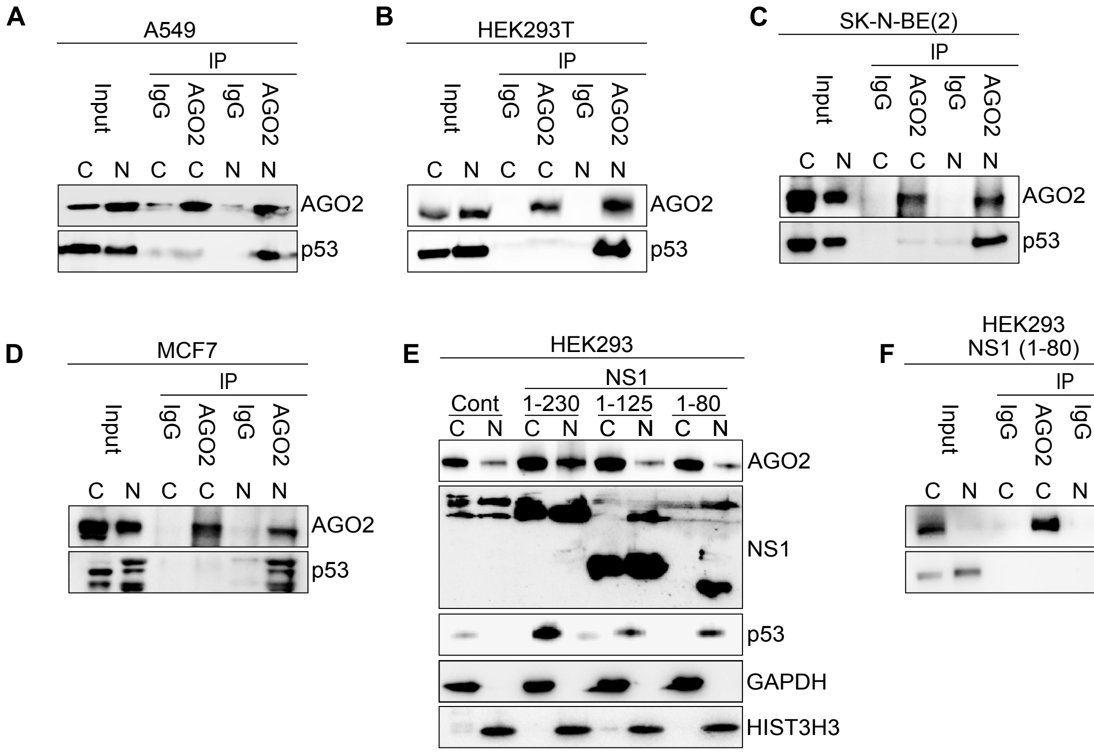


**Supplemental figure 2, related to main figure 2**

**(A)** AGO2 immunoprecipitation (IP) from cytoplasmic or nuclear fraction of A549 cells. Representative immunoblots of AGO2 and p53. n= 3

**(B)** same as in **(A)** but in HEK293T cells.

**(C)** same as in **(A)** but in SK-N-BE(2) cells.

**(D)** same as in **(A)** but in MCF7 cells.

**(E)** Representative AGO2, NS1 and p53 immunoblots from cytoplasmic (C) and nuclear (N) lysates in HEK293 cells transfected with WT (1-230) and deletion mutant NS1 (1-80 and 1-124) expressing plasmid for 2 days. GAPDH served as cytoplasmic marker and HIST3H3 as nuclear marker. n=3

**(F)** AGO2 immunoprecipitation (IP) from cytoplasmic or nuclear fraction of HEK293 cells transfected with mutant NS1 expressing plasmids for 24 hours. Representative immunoblots of AGO2 and p53. n= 3

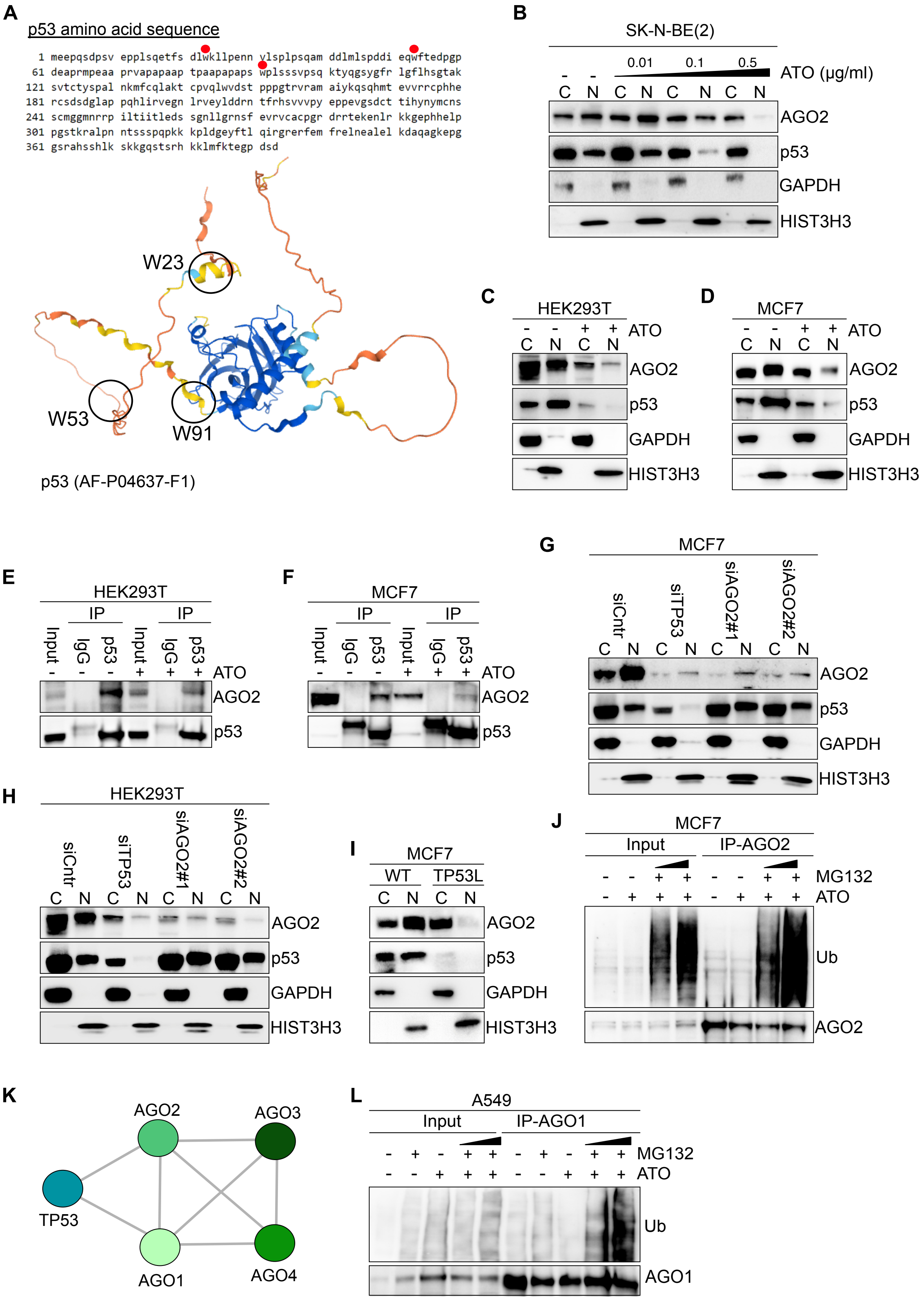


**Supplemental figure 3, related to main figure 3**

**(A)** Sequence and ribbon diagram of predicted human p53 protein structure from the AlphaFold protein structure database (AF-P04637-F1). W23, W53 and W91 are shown in the N-terminal domain and as red dots on the protein sequence.

**(B)** Representative AGO2 and p53 immunoblots from cytoplasmic (C) and nuclear (N) lysates in SK-N-BE(2) cells treated with 0.01, 0.1 or 0.5 µg/mL arsenic trioxide (ATO) for 24 hours. GAPDH served as a cytoplasmic marker and HIST3H3 served as nuclear marker. n= 3

**(C)** Representative AGO2 and p53 immunoblots from cytoplasmic (C) and nuclear (N) lysates in HEK293T cells treated with 0.5 µg/mL arsenic trioxide (ATO) for 24 hours. GAPDH served as a cytoplasmic marker and HIST3H3 served as nuclear marker. n=3

**(D)** Same as in **(C)** but for MCF7 cells.

**(E)** p53 immunoprecipitation (IP) from HEK293T cells treated with 0.5 µg/mL Arsenite trioxide (ATO) for 24 hours. Representative immunoblots of AGO2 and p53. n=3

**(F)** Same as in **(E)** but for MCF7 cells.

**(G)** Representative AGO2 and p53 immunoblots from cytoplasmic (C) and nuclear (N) lysates in MCF7 cells treated with siRNAs for 24 hours. siCntr: control scramble siRNAs; siTP53: siRNA specific for TP53; and siAGO2: two different siRNAs specific for AGO2. GAPDH served as a cytoplasmic marker and HIST3H3 served as nuclear marker. n=3

**(H)** Same as in **(G)** but for HEK293T cells.

**(I)** Representative AGO2 and p53 immunoblots from cytoplasmic (C) and nuclear (N) lysates in WT and TP53L MCF7 cells. GAPDH served as a cytoplasmic marker and HIST3H3 served as nuclear marker. n=3

**(J)** AGO2 immunoprecipitation (IP) from MCF7 cells treated with 2 µg/ml MG132 for 2 hours before additional 24 hours of treatment with 0.5 µg/ml arsenic trioxide (ATO). Representative immunoblots of ubiquitin (Ub) and AGO2. n=3

**(K)** STRING protein-protein interaction network of AGO1, AGO2, AGO3, AGO4 and p53.

**(L)** AGO1 immunoprecipitation (IP) from A549 cells treated with 2 µg/ml MG132 for 2 hours before additional 24 hours of treatment with 0.5 µg/ml arsenic trioxide (ATO). Representative immunoblots of ubiquitin (Ub) and AGO1. n=3

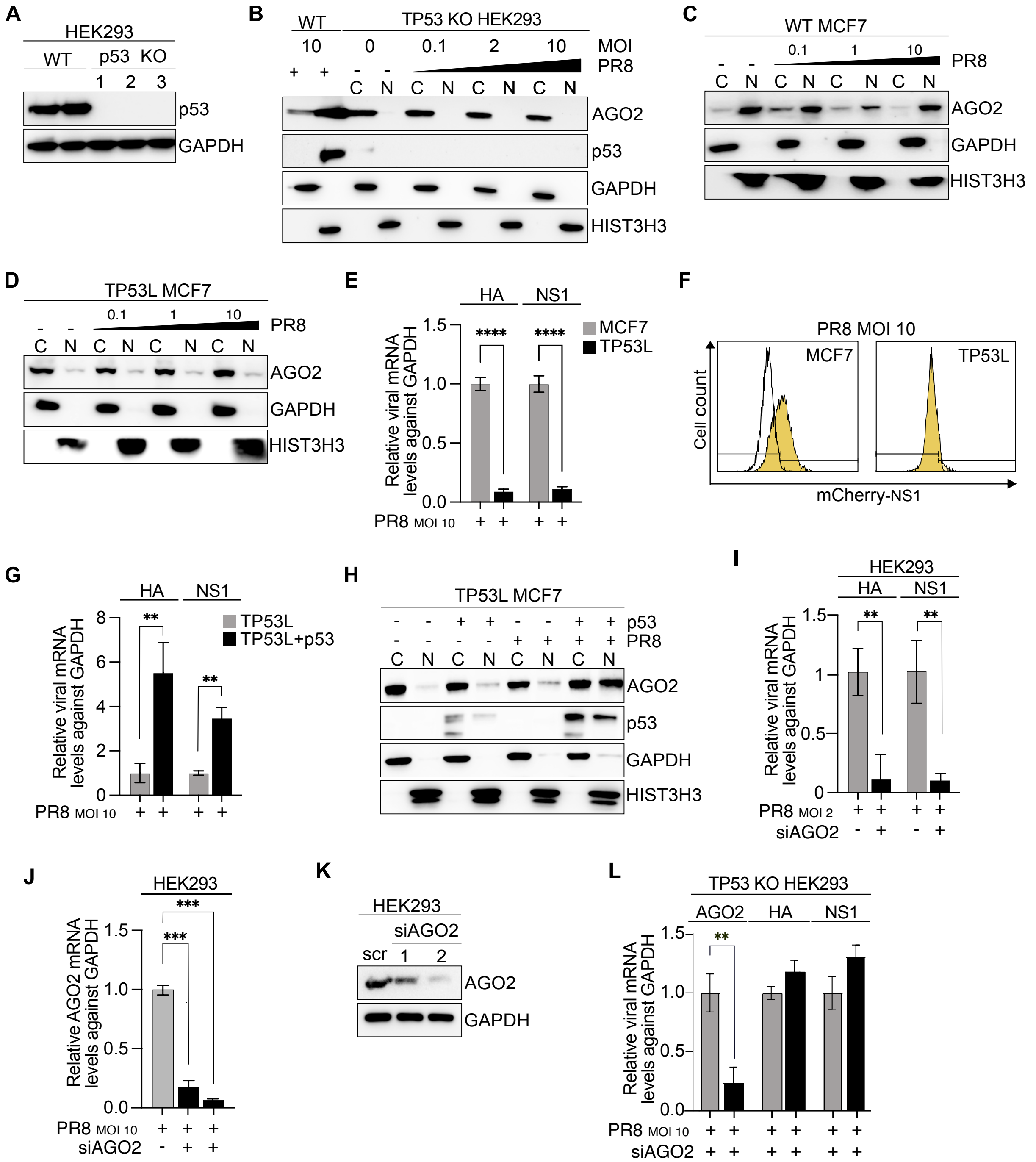


**Supplemental figure 4, related to main figure 4**

**(A)** Representative p53 immunoblots from whole cell lysates of WT and TP53 KO HEK293 cells. There individual KO clones are indicated. GAPDH served as a loading control. n=1

**(B)** Representative AGO2 and p53 immunoblots from cytoplasmic (C) and nuclear (N) lysates in WT and TP53 KO HEK293 cells infected with PR8 virus at MOI 0.1; 2; 10 for 16 hours. GAPDH served as a cytoplasmic marker and HIST3H3 served as nuclear marker. n=3

**(C)** Representative AGO2 immunoblots from cytoplasmic (C) and nuclear (N) lysates in WT MCF7 cells infected with PR8 virus at MOI 0.1; 1; 10 for 16 hours. GAPDH served as a cytoplasmic marker and HIST3H3 served as nuclear marker. n=3

**(D)** Same as in **(C)** but in TP53L MCF7 cells.

**(E)** Relative expression, as measured by RT-qPCR, of NS1 and HA mRNA levels in WT and TP53L MCF7 cells upon infection with PR8 virus at MOI 10 for 16 hours. GAPDH was used as a reference gene. Bars are mean and error bars represent ± SD. **** $p<0.0001$ by unpaired t-test. n=3

**(F)** Flow cytometry analysis of mCherry expression in WT and TP53L MCF7 cells upon infection with PR8-mCherry virus at MOI 10 for 16 hours. White histogram shows mock-infected cells while yellow histogram is PR8-infected. n=3

**(G)** Relative expression, as measured by RT-qPCR, of NS1 and HA mRNA levels in WT TP53L MCF7 cells. TP53L MCF7 cells were transfected with Flag-WT-p53 expressing plasmids for 24 hours. Subsequently, both WT and TP53L MCF7 cells overexpressing Flag-WT-p53 transiently were infected with PR8 virus at MOI 10 for 16 additional hours. GAPDH was used as a reference gene. Bars are mean and error bars represent ± SD. ** $p<0.01$ by unpaired t-test. n=3

**(H)** Representative AGO2 and p53 immunoblots from cytoplasmic (C) and nuclear (N) lysates in TP53L MCF7 cells transfected with Flag-WT-p53 expressing plasmids for 24 hours and infected with PR8 virus at MOI 10 for 16 additional hours. GAPDH served as a cytoplasmic marker and HIST3H3 served as nuclear marker. n=3

**(I)** Relative expression, as measured by RT-qPCR, of HA and NS1 mRNA levels in HEK293 cells treated with siRNAs against AGO2 for 24 hours. 16 hours before the end of incubation, cells were infected with PR8 virus at MOI 10. GAPDH was used as a reference gene. Bars are mean and error bars represent ± SD. *** $p<0.001$ by unpaired t-test. n=3

**(J)** Relative expression, as measured by RT-qPCR, of AGO2 mRNA levels in HEK293 cells treated with siRNAs against TP53 or AGO2 for 48 hours. 16 hours before the end of incubation, cells were infected with PR8 virus at MOI 10. GAPDH was used as a reference gene. Bars are mean and error bars represent ± SD. *** $p<0.001$ by unpaired t-test. n=3

**(K)** Representative AGO2 immunoblots in HEK293 treated with siRNAs against AGO2 for 24 hours.

**(L)** Relative expression, as measured by RT-qPCR, of AGO2, NS1 and HA mRNA levels in TP53 KO HEK293 cells treated with siRNAs against AGO2 for 24 hours. 16 hours before the end of incubation, cells were infected with PR8 virus at MOI 10. GAPDH was used as a reference gene. Bars are mean and error bars represent ± SD. ** $p<0.01$ by unpaired t-test. n=3

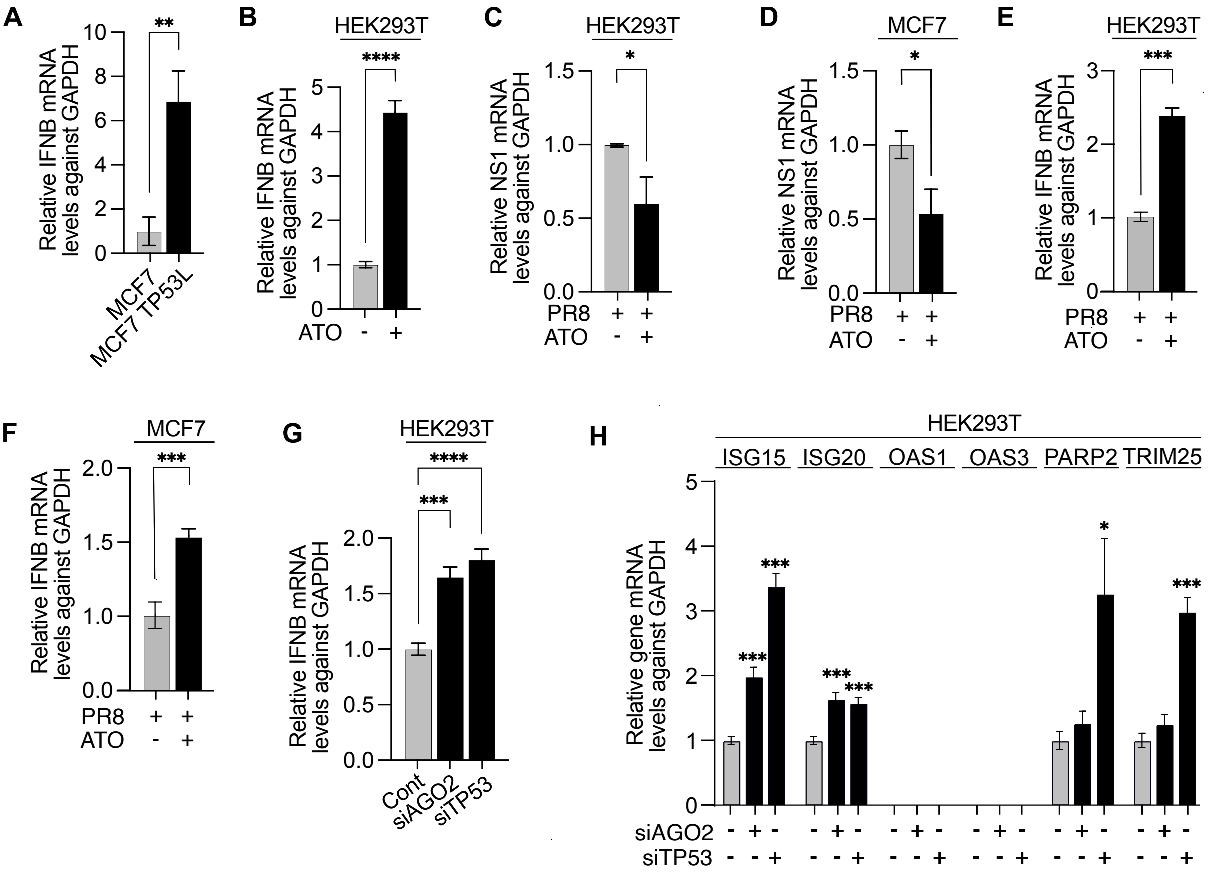


**Supplemental figure 5, related to main figure 5**

**(A)** Relative expression, as measured by RT-qPCR, of IFNB mRNA levels in WT and TP53L MCF7 cells. GAPDH was used as a reference gene. Bars are mean and error bars represent ± SD. ** $p<0.01$ by unpaired t-test. n=3

**(B)** Relative expression, as measured by RT-qPCR, of IFNB mRNA levels in HEK293T cells treated for 24 hours with 0.5 µg/ml arsenic trioxide (ATO) or vehicle. GAPDH was used as a reference gene. Bars are mean and error bars represent ± SD. **** $p<0.0001$ by unpaired t-test. n=3

**(C)** Relative expression, as measured by RT-qPCR, of NS1 mRNA levels in HEK293T cells treated for 24 hours with 0.5 µg/ml arsenic trioxide (ATO) or vehicle and infected with PR8 virus at MOI 10 for 16 hours. GAPDH was used as a reference gene. Bars are mean and error bars represent ± SD. * $p<0.05$ by unpaired t-test. n=3

**(D)** Same as in **(C)** but in MCF7 cells * $p<0.05$ by unpaired t-test. n=3

**(E)** Relative expression, as measured by RT-qPCR, of INFB mRNA levels in HEK293T cells treated for 24 hours with 0.5 µg/ml arsenic trioxide (ATO) or vehicle and infected with PR8 virus at MOI 10 for 16 hours. GAPDH was used as a reference gene. Bars are mean and error bars represent ± SD. *** $p<0.001$ by unpaired t-test. n=3

**(F)** same as in **(E)** but in MCF7 cells. *** $p<0.001$ by unpaired t-test. n=3

**(G)** Relative expression, as measured by RT-qPCR, of IFNB mRNA levels in HEK293T cells treated with siRNAs against TP53 or AGO2 for 48 hours. GAPDH was used as a reference gene. Bars are mean and error bars represent ± SD. *** $p<0.001$, **** $p<0.0001$ by unpaired t-test. n=3

**(H)** Normalized luciferase signal of ISRE-transfected HEK293T cells, treated with siRNA against AGO2 and TP53 for 48 hours. 16 hours before the end of incubation, cells were infected with PR8 virus at MOI 10. GAPDH was used as a reference gene. Bars are mean and error bars represent ± SD. * $p<0.05$, *** $p<0.001$ by unpaired t-test. n=3

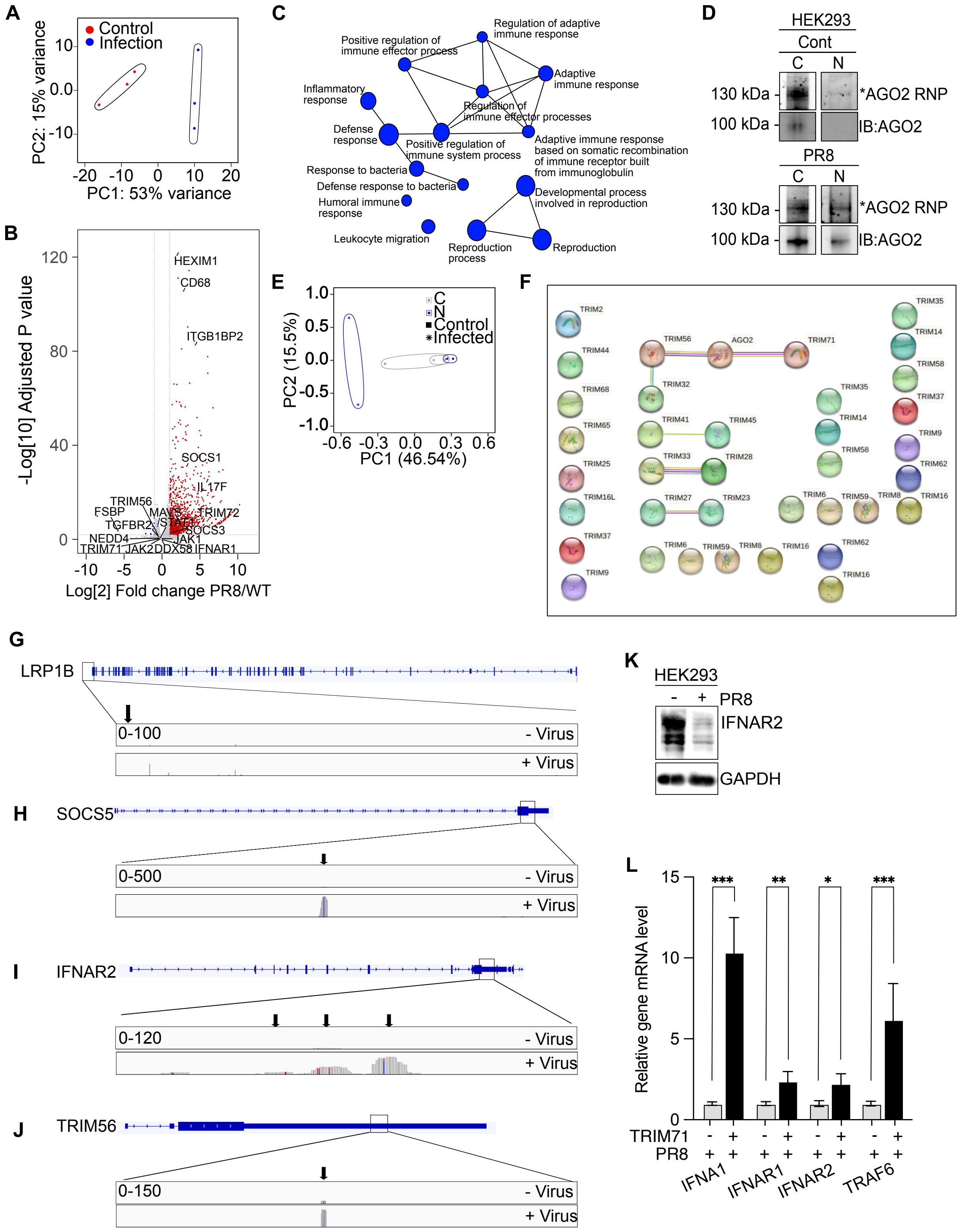


**Supplemental figure 6, related to main figure 6**

**(A)** Principal component analysis (PCA) plot of RNAseq from HEK293 cells infected for 16 hours with PR8 at MOI 10 (blue) or mock infected (red).

**(B)** Volcano plot showing RNAseq results of differentially expressed genes between PR8 infected and non-infected HEK293 cells. In red are the upregulated genes while in blue the downregulated.

**(C)** Top 15 significantly upregulated Gene Ontology (GO) pathways based on genes upregulated during PR8 infection in HEK293 cells.

**(D)** IR680 fluorescent image of crosslinked and fluorescent adapter-ligated AGO1-4 ribonucleoprotein complexes from the cytoplasmic (C) and nuclear (N) fractions of HEK293 cells with or without PR8 viral infection, separated by SDS-PAGE. AGO2 immunoblots depict the efficiency of the pulldown. n=2

**(E)** Principal component analysis (PCA) plot of AGO-fPAR-CLIP from cytoplasmic (C, grey) and nuclear (N, blue) fractions of HEK293 cells infected for 16 hours with PR8 at MOI 10 (star) or mock infected (square).

**(F)** STRING protein-protein interaction network of AGO2 and TRIM71 (or TRIM family proteins).

**(G)** LRP1B IGV track in nuclear fraction of control (- Virus) and IAV infected (+Virus) HEK293 cells.

**(H)** same as in **(G)** but for SOCS5.

**(I)** same as in **(G)** but for IFNAR2.

**(J)** same as in **(G)** but for TRIM56

**(K)** Representative IFNAR2 immunoblot from whole cell lysates in HEK293 infected with PR8 virus at MOI 10 for 16 hours. GAPDH served as a loading control. n= 3

**(L)** Relative expression, as measured by RT-qPCR, of IFNA, IFNB, IFNAR1, IFNAR2 and TRAF6 mRNA levels in HEK293 cells transfected with HA-TRIM71 and infected with PR8 virus at MOI 10 for 16 hours. GAPDH was used as a reference gene. Bars are mean and error bars represent ± SD. ∗ $p<0.05$, ∗∗ $p<0.01$, ∗∗∗ $p<0.001$ by unpaired t-test. n= 3

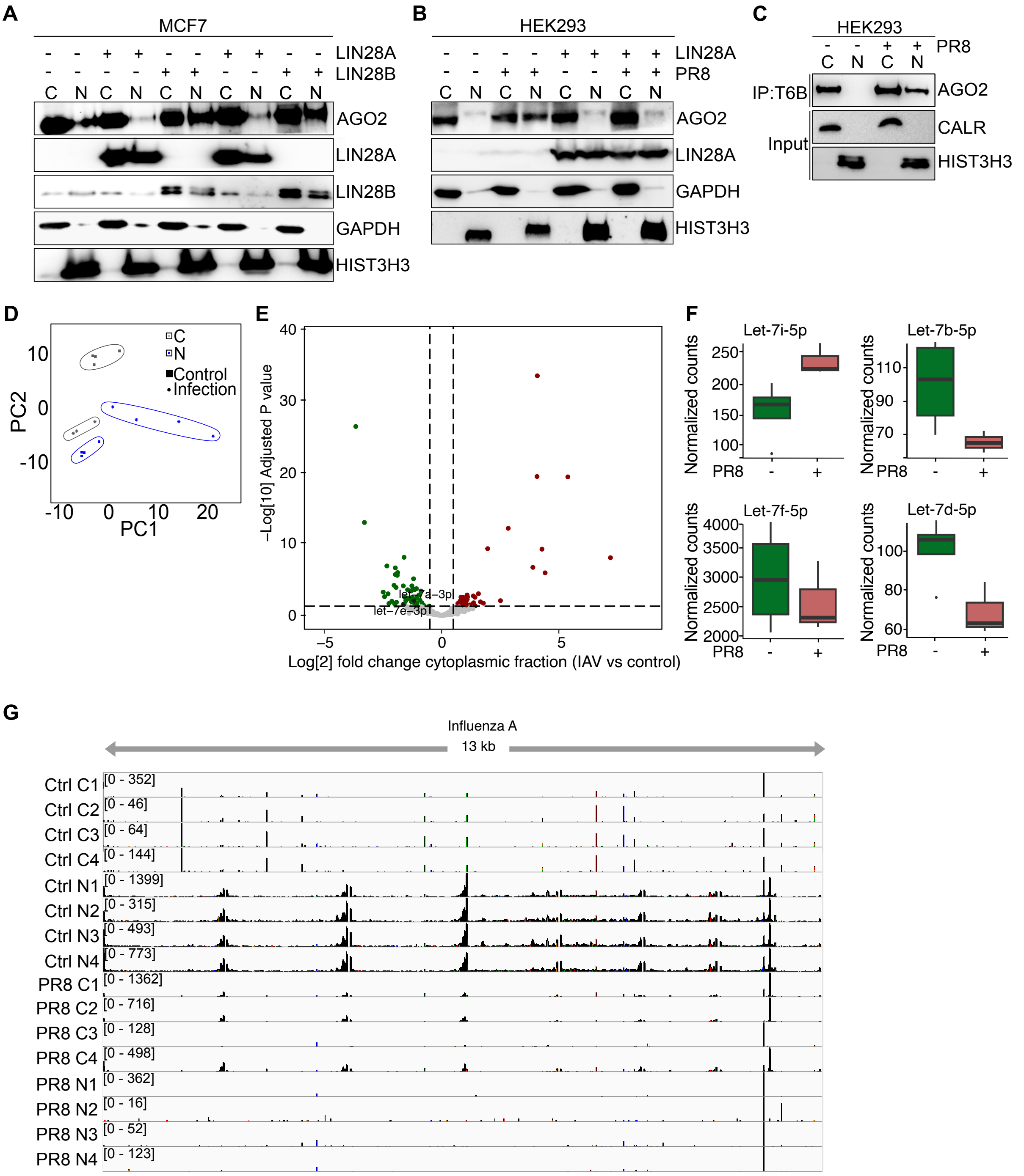


### Supplemental figure 7, related to main figure 7

**(A)** Representative AGO2, LIN28A and LIN28B immunoblots from cytoplasmic (C) and nuclear (N) lysates from MCF7 cells transfected with V5-LIN28A or Flag-LIN28B expressing plasmids for 24 hours. GAPDH served as a cytoplasmic marker and HIST3H3 served as nuclear marker. n=2

**(B)** Representative AGO2 and LIN28A immunoblots from cytoplasmic (C) and nuclear (N) lysates from HEK293 cells transfected with V5-LIN28A expressing plasmids for 24 hours. 16 hours before the end of incubation, cells were infected with PR8 virus at MOI 10. GAPDH served as a cytoplasmic marker and HIST3H3 served as nuclear marker. n=2

**(C)** AGO1-4 immunoprecipitation (IP) using T6B peptide from cytoplasmic (C) and nuclear (N) lysates of HEK293 cells. Cells were infected with PR8 virus at MOI 10 for 16 hours. Representative immunoblots of AGO2. Calreticulin (CALR) served as a cytoplasmic marker and HIST3H3 served as nuclear marker n= 4

**(D)** Principal component analysis (PCA) plot of AGO-miRNAseq from cytoplasmic (C, grey) and nuclear (N, blue) fractions of HEK293 cells infected for 16 hours with PR8 at MOI 10 (star) or mock infected (square).

**(E)** Volcano plot showing miRNAseq results of differentially expressed miRNAs from the cytoplasmic fraction of PR8 infected and non-infected HEK293 cells. In red are the upregulated genes while in green the downregulated.

**(F)** Significantly differentially expressed Let7i/b/f/d-5p from the cytoplasmic fraction of PR8 infected and non-infected HEK293 cells.

**(G)** IGV track in cytoplasmic and nuclear fraction of control and PR8-IAV infected HEK293 cells mapped to the IAV genome.

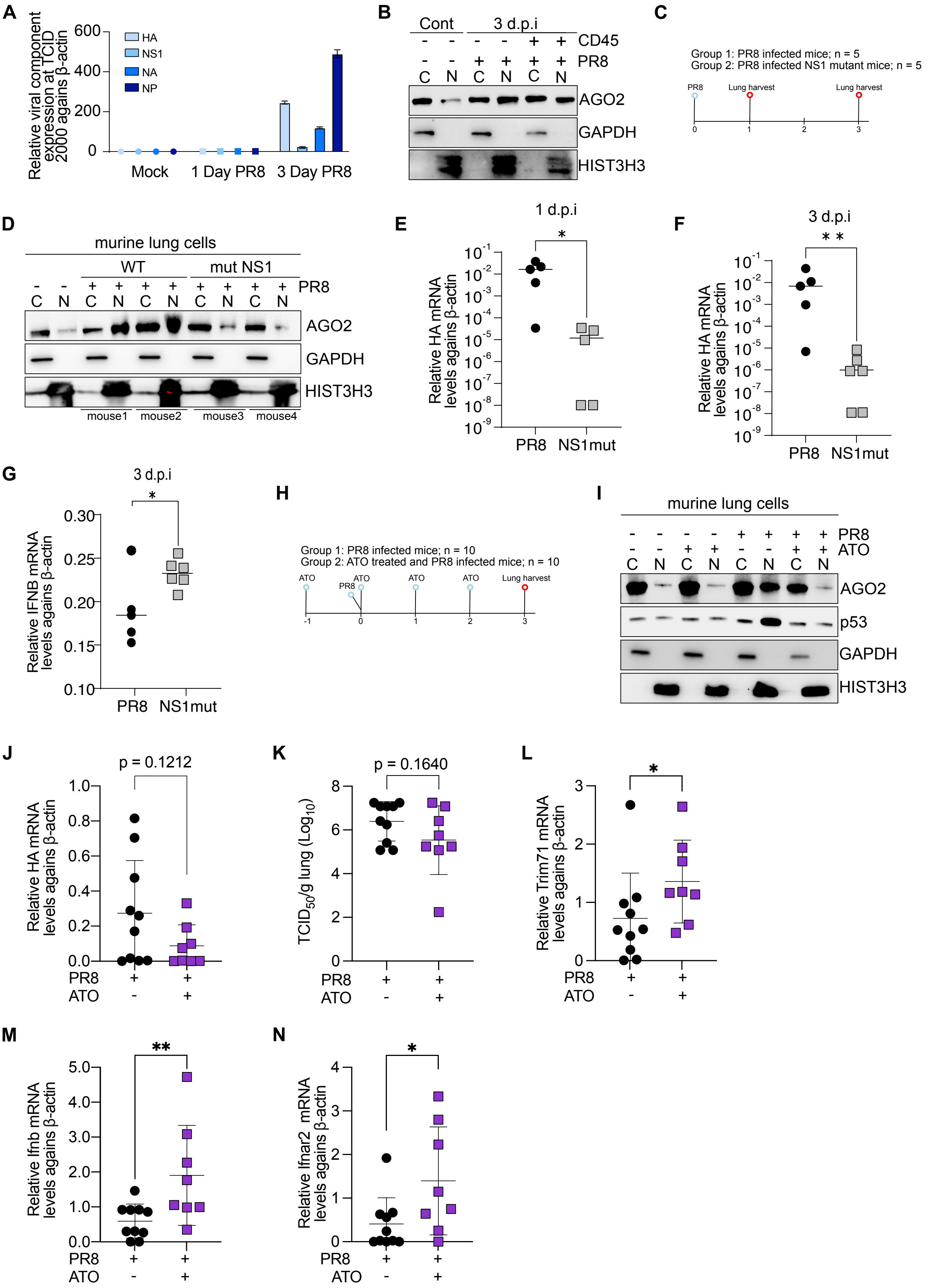


**Supplemental figure 8, related to main figure 8**

**(A)** Relative expression, as measured by RT-qPCR, of NS1, HA, NA and NP mRNA levels in lung cells isolated from mice upon infection with PR8 virus at TCID50 2000 for 1 or 3 days.

β-actin was used as a reference gene. Bars are mean and error bars represent ± SD. n= 2 independent experiments with 2 mice each

**(B)** Representative AGO2 immunoblots from cytoplasmic (C) and nuclear (N) lysates in lung cells isolated from mice at 3 days post infection. Shown are both the $CD45^+$ (immune cells) and $CD45^-$ (stromal cells) fractions. GAPDH served as a cytoplasmic marker and HIST3H3 served as nuclear marker. n= 2 independent experiments with 2 mice each.

**(C)** Schematic representation of the experimental setup for experiments in **(D-G)**. Mice were infected i.n. with 250 $TCID_{50}$ of PR8 or $NS1_{1-124}$ mutant and lungs harvested at 1 or 3 days post-infection.

**(D)** Representative AGO2 immunoblots from cytoplasmic (C) and nuclear (N) lysates in lung cells isolated from mice infected with PR8 or $NS1_{1-124}$ mutant. GAPDH served as a cytoplasmic marker and HIST3H3 served as nuclear marker. n=1 independent experiments with 5 mice each.

**(E)** Graph representing HA mRNA levels at 1 day post infection in lung cells isolated from mice infected with PR8 or $NS1_{1-124}$ mutant. β-actin was used as a reference gene. Bars are mean and error bars represent ± SD. n=1 independent experiments with 5 mice each. ∗ $p<0.05$ by unpaired t-test.

**(F)** same as in **(E)** but for 3 days post infection. ∗∗ $p<0.01$by unpaired t-test.

**(G)** same as in **(F)** but for INFB. ∗ $p<0.05$ by unpaired t-test

**(H)** Schematic representation of the experimental setup for experiments in **(I-N)**. Mice were treated daily with i.p. injections of 0.15 mg/kg ATO in PBS (or vehicle), starting from day -1. At day 0 mice were infected i.n. with 2000 $TCID_{50}$ PR8 and lungs harvested at 3 days post-infection.

**(I)** Representative AGO2 and p53 immunoblots from cytoplasmic (C) and nuclear (N) lysates in lung cells isolated from mice treated with ATO and PR8 infected. GAPDH served as a cytoplasmic marker and HIST3H3 served as nuclear marker. n=3 independent experiments.

**(J)** Graph representing HA mRNA levels in lung cells isolated from mice treated with ATO (or vehicle) and PR8 infected. Shown are the individual mice with bar representing the mean and error bars represent ± SD. Statistical analysis was performed by unpaired t-test. n=3 independent experiments with a total of 10 (PR8) and 8 (PR8-ATO) mice.

**(K)** Graph representing $\log_{10}$ $TCID_{50}$/g lung in mice treated with ATO (or vehicle) and PR8 infected. Shown are the individual mice with bar representing the mean and error bars represent ± SD. Statistical analysis was performed by unpaired t-test. n=3 independent experiments with a total of 10 (PR8) and 8 (PR8-ATO) mice.

**(L)** Graph representing the mRNA levels of *Trim71* in lung cells isolated from mice treated with ATO (or vehicle) and PR8 infected. Shown are the individual mice with bar representing the mean and error bars represent ± SD. * $p<0.05$ by unpaired t-test. n=3 independent experiments with a total of 10 (PR8) and 8 (PR8-ATO) mice.

**(M)** same as in **(L)** but for *Ifnb*. ∗∗ $p<0.01$ by unpaired t-test.

**(N)** same as in **(L)** but for *Ifnar2*. ∗ $p<0.05$ by unpaired t-test.
